# Evaluation of Active Learning Selection Strategies and Characterization of Informative Sequences for Sequence-to-Expression Models

**DOI:** 10.64898/2026.05.21.727038

**Authors:** Justin Qian, Abdul Muntakim Rafi, Emmanuel Cazottes, Carl G. de Boer

## Abstract

DNA sequence-to-expression models have advanced rapidly, yet they still generalize poorly beyond their training distribution, limiting their use for tasks such as variant effect prediction. Active learning has improved data efficiency across many machine learning domains, but no large-scale study has benchmarked selection strategies for sequence-to-expression models using real experimental data or characterized the sequences they select. We benchmarked six active learning strategies across diverse model architectures, datasets, and configurations. All strategies outperformed random sampling, with uncertainty-based methods performing best. Most of the gains achievable through many small acquisition rounds could be matched with fewer, larger rounds, making lab-in-the-loop workflows experimentally practical. Different strategies selected substantially overlapping sets of sequences that occupied distinct regions of sequence space and were enriched for higher expression, specific dinucleotide compositions, and denser transcription factor binding sites. Nevertheless, active learning consistently outperformed selection based on these biological properties alone, indicating that informativeness is not fully captured by any single feature. Together, our results establish active learning as a critical tool for improving sequence-to-expression models, identify biological signatures of informative sequences, and lay the foundation for iterative lab-in-the-loop refinement.

## Introduction

Deep learning models have made substantial improvements in deciphering cis-regulatory logic^1–7^. While these sequence-to-expression approaches have demonstrated progress driven by advances in model architectures^8–11^, training strategies^12–14^, and, most importantly, dataset expansion^15–17^, they face significant challenges regarding generalization^18–24^. A primary limitation of current dataset expansion approaches is that genomic sequences represent only a tiny fraction of the theoretical sequence space^25,26^. While modern synthesis technologies allow for the creation of massive sequence libraries to augment the limited genomic training data^27–31^, experimental profiling remains costly and time-consuming^32^. This raises a fundamental question: which sequences should be profiled to maximize improvements in model performance and mechanistic understanding?

Active Learning (AL), a machine learning method where the model selects the most informative data from an unlabeled pool for labeling^33^, offers a potential solution. AL is premised on the hypothesis that after training an initial model on a set of available data, the model can be used to identify the most informative data for subsequent model refinement, by focusing on novel examples while ignoring redundant or previously learned instances. This can help direct limited experimental resources towards characterizing the most informative samples, and consequently, maximizing model performance. AL was originally used in classification tasks by classical machine learning algorithms^33^, and has been expanded to regression tasks and deep neural networks^34–36^. Many AL strategies have involved selecting samples based on the degree of uncertainty in model predictions^33,37–39^, and numerous uncertainty-based algorithms have been developed that are compatible with regression tasks such as DNA sequence-to-expression prediction^40–43^. However, selecting a batch of the most uncertain examples may result in significant redundancy within batches^36^, which led to the development of diversity-focused strategies. Diversity-based methods sacrifice exploitation of the most uncertain examples but avoid redundancy, making them a popular choice in AL^44,45^.

DNA sequences are an ideal setting for AL since any sequence can be synthesized and tested in a high-throughput fashion using Massively Parallel Reporter Assays (MPRAs), enabling exploration of a vast sequence space. This makes sequence-to-expression modeling a natural setting for lab-in-the-loop experimentation, where model predictions guide sequence design and selection, experimental assays generate new measurements, and the resulting data are fed back to iteratively improve the model. Indeed, several recent studies have demonstrated the potential of AL to improve sequence-to-expression models. Friedman et al.^46^ used AL with MPRAs to study the photoreceptor transcription factor cone-rod homeobox (CRX) in mouse retina, demonstrating that AL can uncover context-dependent regulatory logic not learnable from genomic sequences alone. Crnjar et al.^47^ developed PIONEER, a computational framework using in silico oracles to systematically benchmark sequence generation strategies and acquisition functions, evaluating model performance under varying distributional shifts. Morrow et al.^48^ applied AL to optimize 3’ UTRs for mRNA therapeutics, achieving 2-fold half-life increases *in vitro* and 30-100-fold protein improvements in mouse models. Shen et al.^49^ compared AL to one-shot optimization for yeast promoter design, showing that AL outperforms traditional approaches in complex epistatic landscapes. Furthermore, Yin et al.^50^ showed that small, targeted additions to training data had the potential to greatly improve model performance in their study on synthetic enhancer design. Critical questions remain about (1) how different AL selection methods compare to each other and to random sampling, (2) the impact of the number of sequences selected per round of AL, (3) how performance varies across biological systems with different scales and regulatory complexity, and perhaps most importantly, (4) what features make AL-selected sequences more informative than others.

In this study, we developed nextFrag, a comprehensive AL framework for benchmarking sequence selection strategies across model architectures, datasets, and experimental configurations, and for characterizing the biological properties that drive model improvement. We benchmarked six AL sequence selection methods using three neural network architectures across two large-scale sequence-to-expression datasets differing in species, scale, and sequence library composition. Across in-distribution (ID), variant effect prediction (VEP), and out-of-distribution (OOD) test sets, AL delivered major performance gains. We examined how a single larger round of AL compares to multiple smaller iterative rounds, showing that the former captures most of the performance gains of the latter. Different AL strategies converged on overlapping sets of sequences, concentrated in specific regions of sequence space. We characterized those sequences and found that AL-selected sequences differ in their expression levels, *in silico mutagenesis* (ISM) scores, sequence composition, and transcription factor binding site (TFBS) occurrences. Finally, when compared to acquisition functions based purely on those simple biological features, AL selection strategies were consistently the most reliable at producing large performance gains.

## Results

### AL benchmarking workflow

To benchmark AL strategies for sequence-to-expression modeling, we simulated iterative selection experiments on real datasets. We trained models with subsets of each dataset, and then used AL approaches to select the most informative sequences from the remaining pool. This approach allowed us to simulate the AL process without requiring new experiments. To maximize the available sequence pool sizes, we focused on two of the largest sequence-to-expression datasets: the Gosai et al. K562 dataset^51^ (∼370,000 reference allele sequences; "human") and the Random Promoter DREAM Challenge dataset^52^ (∼6,700,000 random sequences; "yeast") (**Fig. 1**). While both datasets reflect experimental measurements of DNA sequences using MPRAs, they differ substantially in sequence library composition: the human dataset consists of genomic sequences, whereas the yeast dataset consists of random sequences.

**Figure 1:**
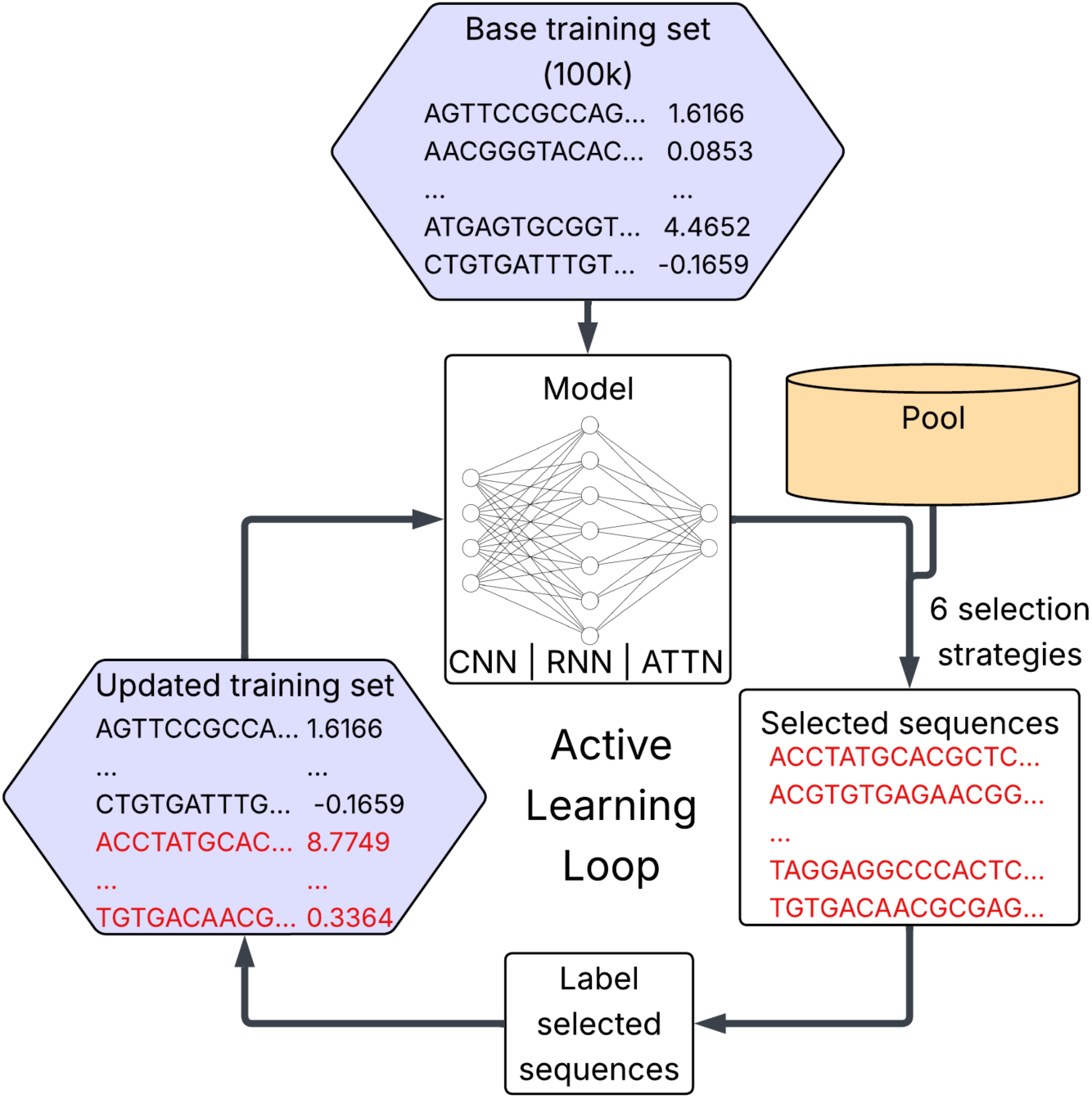
Overview of AL workflow. Neural network models (RNN, CNN, ATTN) are trained on 100,000 sequences. Models select sequences from the pool using one of six selection strategies. These sequences are then labeled and added to the training set, and the model is retrained, and the process is repeated.

Our AL benchmarks used the state-of-the-art DREAM Challenge MPRA model architectures^52^, selected for their high performance across diverse sequence-to-expression tasks and spanning a variety of architecture types (RNN, CNN, ATTN). Each model was initially trained on a base training set of 100,000 sequences, randomly sampled from each dataset. The limited training data left most sequences untouched, maximizing the selection pool sizes while still resulting in decent baseline model performance. We then performed AL using six selection strategies, categorized as uncertainty-based and diversity-based. Uncertainty-based strategies, which prioritize sequences where model predictions are least consistent, included Monte Carlo (MC) dropout, which applies dropout at inference time to make multiple predictions with a single model, and three types of ensembles differing in composition and size: (1) “all arch” ensemble of all three architectures, (2) “RNN+CNN” ensemble of the two best performing architectures, (3) single-architecture ensembles (“RNN ensemble”, the best performing, shown in main results, and “CNN ensemble”/“ATTN ensemble” shown in supplement). Multi-architecture ensembles identify informative sequences by leveraging disagreements between different model architectures, whereas single-architecture ensembles and MC dropout identify sequences where even the same model architecture, including the strongest, produces inconsistent predictions. Diversity-based strategies focus on selecting a broad range of sequences and minimizing redundant sequence features. We tested k-means clustering, which selects a representative set of sequences from the model’s latent space, and Largest Cluster Maximum Distance (LCMD), which prioritizes maximal latent space distance between selections (see **Methods**). We compared two AL iteration configurations: 3 rounds selecting 20,000 sequences per round, and 1 round selecting 60,000 sequences. In each round, we added the selected sequences to the training set, and then retrained models from scratch instead of fine-tuning to mitigate the risk of catastrophic forgetting^53^ or other consequences from sub-optimal fine-tuning procedures. Performance was compared to a random sampling baseline, selecting the same number of sequences per round.

### AL strategies improve model performance and generalization

We first evaluated the performance of AL strategies against random sampling on a held-out in-domain (ID) test set. Uncertainty-based AL strategies consistently outperformed random sampling after each round in both datasets, achieving in a single round what random sampling required three rounds to match (**Fig. 2A-B**). Diversity-based strategies performed more variably, consistently trailing uncertainty-based strategies but still matching or exceeding the random baseline (**Fig. 2A-B**). Final-round performance distributions across both datasets confirmed that while no single uncertainty-based strategy was universally optimal, they collectively provided superior and more stable gains compared to diversity-based or random selection (**Fig. 2C-D**).

**Figure 2:**
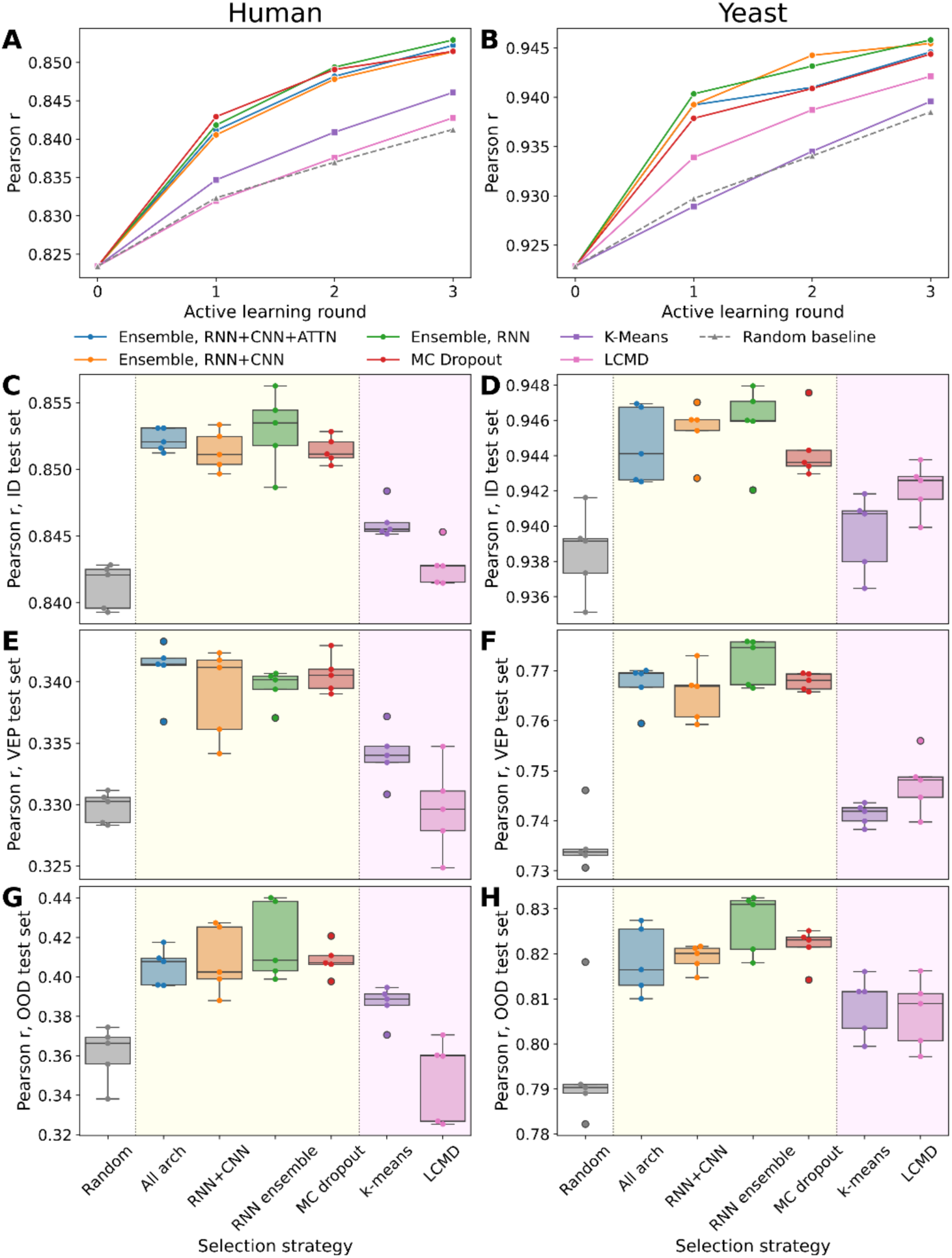
AL outperforms random sampling across test sets. **A-B)** Performance of RNN models (Pearson *r*, *y*-axes) across AL rounds (*x*-axes) for uncertainty-based (solid lines, circular markers) and diversity-based (solid lines, square markers) selection strategies compared to a random sampling baseline (dashed gray line) on the ID test set, averaged across 5 replicates of RNN for human (**A**) and yeast (**B**) datasets. **C-H)** Performance of RNN models (Pearson *r*, *y*-axes) across AL selection methods (*x*-axes; yellow background for uncertainty- and magenta background for diversity-based strategies) compared to a random sampling baseline (blank background) on the ID test set **(C-D)**, VEP test set **(E-F)**, and OOD test set **(G-H)**, for human **(C, E, G)** and yeast **(D, F, H)** datasets.

Beyond ID performance, we investigated whether AL improves model generalization, a major challenge in genomics modeling^19,20^, by introducing two benchmarks on distinct tasks. First, we evaluated performance on the variant effect prediction (VEP) task. In the human dataset, this comprised expression differences between reference and alternate alleles for genomic test set sequences. In yeast, it consisted of pairs of sequences, either random or genomic, differing by a single nucleotide (the "SNV" test subset from the DREAM Challenge^52^). Second, we assessed performance on out-of-domain (OOD) test sets where the sequence origin differed from the training data: synthetic sequences for human models, genomic sequences for yeast models. Uncertainty-based AL substantially improved performance on these generalization benchmarks (**Fig. 2E-H**). In most cases, these gains exceeded the relative gains observed on ID test data, with an increase in *r^2^* of ∼3% for human OOD and ∼6% for yeast VEP/OOD for some uncertainty strategies, compared to ∼1.5% for ID in both datasets. Notably, diversity-based selection (k-means) improved VEP and OOD performance over random sampling in yeast despite not showing substantial ID gains, suggesting that diversity-based selection can specifically benefit generalization.

These trends were largely consistent between architectures (**Supplementary Fig. 1, 2**), with the only exception being yeast ATTN models. For yeast ATTN, AL only improved performance relative to random selection when sequences were selected with the help of non-ATTN models (all arch, RNN+CNN), where performance gains were large (**Supplementary Fig. 2D, F, H**). Strategies that relied exclusively on the ATTN model (ATTN ensemble, MC dropout, k-means, LCMD) did not see this improvement (**Supplementary Fig. 2D, F, H**). The performance of the initial yeast ATTN model was lower than RNN and CNN (Pearson *r* ∼ 0.89 for ATTN, compared to ∼0.925 for RNN/CNN; **Fig. 2B, Supplementary Fig. 1B, 2B**), suggesting that it may have been data-limited. We reason that this initial deficit produced poor uncertainty estimates, which in turn degraded ATTN-driven acquisition.

Across all three test sets, uncertainty-based strategies consistently outperformed both diversity-based and random selection. To understand what drives this advantage, we examined whether uncertainty-based strategies were in fact selecting sequences the model struggles to predict. We classified sequences into those frequently selected across uncertainty-based strategies ("often-uncertain"; see **Methods**) and those consistently avoided ("rarely-uncertain"; see **Methods**), and calculated how well the models could predict ground truth measurements for each set. Often-uncertain sequences had significantly higher prediction error than the overall AL pool, while rarely-uncertain sequences had significantly lower error (**Supplementary Fig. 3**), confirming that uncertainty serves as an effective proxy for prediction error.

### A single large AL round performs similarly to multiple smaller AL rounds

We next sought to evaluate how the AL round size affected performance. Fewer larger rounds would be more practical for most applications in genomics since sequence annotation experiments (e.g., an MPRA) are time-consuming and the cost of sequence synthesis increases sub-linearly. Hence, if comparable performance can be obtained, fewer rounds would be preferred. To determine whether AL remains effective when iterations are reduced, we compared three sequential rounds of AL selecting 20,000 sequences each (“3x20k”) with a single round selecting the same total number of sequences (“1x60k”).

Model performance was broadly similar whether using three rounds or a single larger round of selection. RNN usually performed slightly better in the 3x20k configuration, but the difference in median *r*^2^ values between configurations in both datasets was, in most cases, less than 0.004 (**Fig. 3A-B**), substantially less than the improvement achieved by uncertainty-based strategies compared to random sampling (0.017 for human, 0.011 for yeast). Similarly, small differences, going in both directions, were observed for CNN and ATTN for the human dataset (**Supplementary Fig. 4A-C**). Only ATTN in yeast showed meaningful differences (**Supplementary Fig. 4D**), but this was also the case where AL did not perform well (**Supplementary Fig. 2D**). Overall, the slight reduction in performance in the 1x60k configuration is consistent with later-round 3x20k models identifying informative sequences more effectively than the initial model, used in the 1x60k configuration for all selections. Nonetheless, this gap is small relative to the gains AL provides over random sampling, indicating that most of the benefit of AL can be captured in a single large selection round.

**Figure 3:**
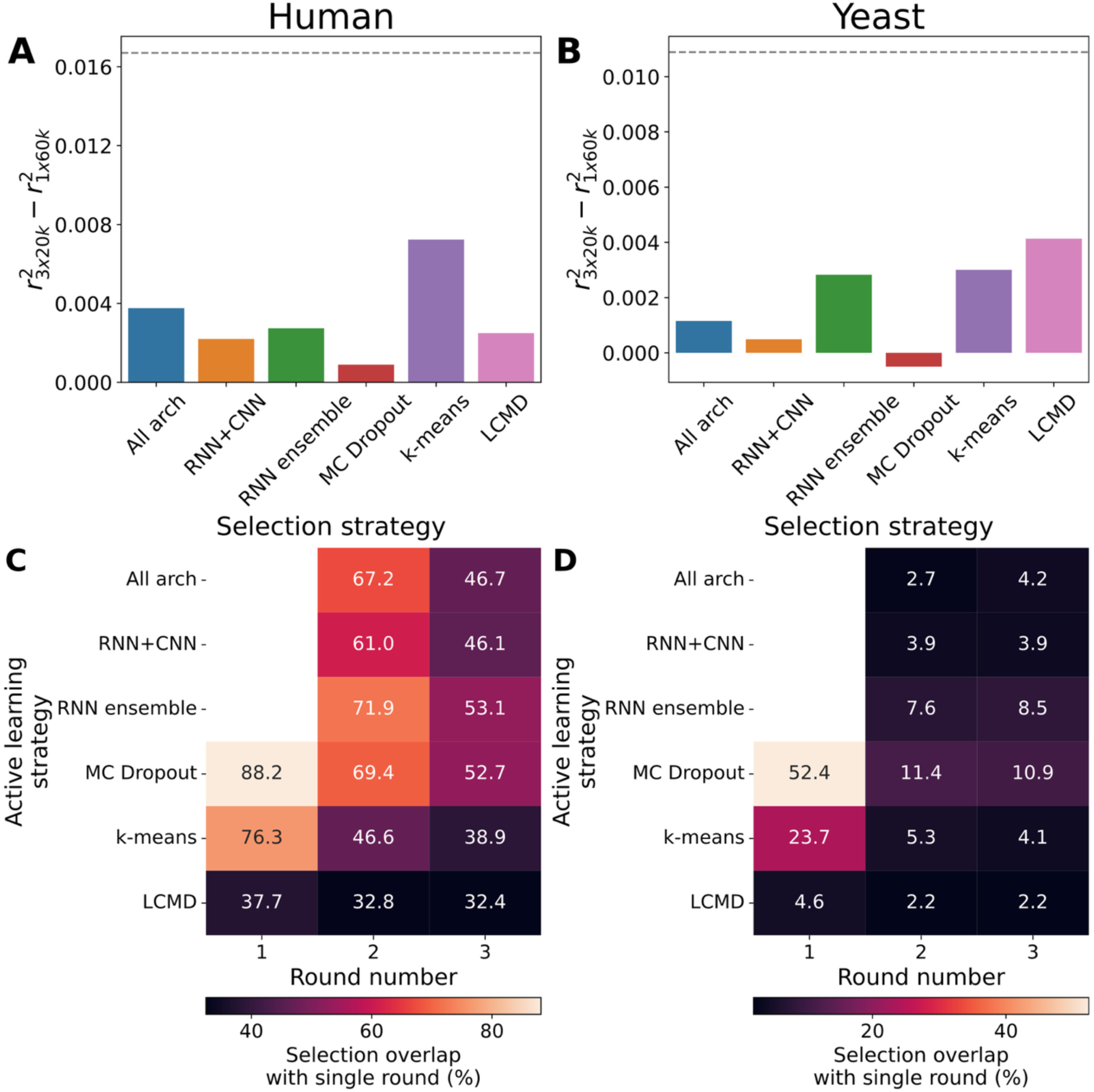
Single-round AL performs comparably compared to three-round AL, even if selected sequences partially differ. **A-B)** Difference in median *r*^2^ between 3x20k and 1x60k configurations for RNN models (*y*-axes), broken down by selection strategy (*x*-axes) in the human (**A**) and yeast (**B**) datasets. The average difference between uncertainty-based strategies and random sampling is indicated by the gray dashed line. **C-D)** Proportion of the 20k sequences selected in each round of the 3x20k configuration (*x*-axes) that are shared with the corresponding 1x60k selection, for each selection strategy (*y*-axes), in the human (**C**) and yeast (**D**) datasets. Round 1 for ensemble methods (All arch, RNN+CNN, RNN ensemble) is not shown since selection is deterministic in that round. All data shown are for RNN.

We next asked whether the similar performance observed between the 3x20k and 1x60k configurations was due to similar sequences being selected. We computed the overlap between sequences selected by the 1x60k configuration and the sequences selected in each round of the 3x20k configuration, with 100% overlap meaning that all 20,000 sequences selected in a given round were also selected in the 1x60k configuration. As expected, first-round selections showed the greatest overlap across all strategies. Because selection is deterministic for ensemble-based strategies, the round 1 selection of the 3x20k configuration corresponds exactly to the top 20k samples chosen by the 1x60k configuration, yielding 100% overlap. (**Fig. 3C, D**). Overlap decreased in subsequent rounds as later-round models diverged in their selections. In human, uncertainty-based strategies retained moderately high overlap even in round 3 (46.7%–53.1%; **Fig. 3C**). Diversity-based strategies showed lower overlap: k-means fell to 38.9% and LCMD to 32.4% by round 3 (**Fig. 3C**). In yeast, the decline was more dramatic, consistent with the much larger pool: by round 2, most strategies shared fewer than 12% of sequences, and by round 3 shared fewer than 5% (**Fig. 3D**). Similar trends were observed in the CNN and ATTN models (**Supplementary Fig. 5**), though ATTN models generally showed lower overlap. Overall, uncertainty-based strategies had higher overlap between 1x60k and 3x20k configurations than diversity-based strategies, though this distinction lessened in later rounds.

### AL strategies display notable, but not complete, overlap in selection

Since AL strategies often achieved similar performance, we next investigated the selection similarities between strategies. To that end, we computed the pairwise overlap of the 60,000 sequences selected by each strategy in the 3x20k configuration, defined as the proportion of sequences selected by both strategies in a given pair and averaged between model replicates.

In the human dataset, uncertainty-based approaches selected more similar subsets (71.0%-75.4% overlap) than expected under random sampling (30.9% overlap) (**Fig. 4A**). Cross-category overlap (uncertainty-based *vs.* diversity-based; 34.7%-50.2%), and overlap within diversity-based approaches were both lower, especially for LCMD whose overlap with all other strategies was the closest to the 30.9% overlap expected by chance (**Fig. 4A**). We observed similar trends in yeast, but the much larger yeast sequence pool compared to human (∼30X) resulted in smaller overlaps overall (usually <20%; **Fig. 4B**). Nonetheless, as with human data, uncertainty-based strategies had a much higher (15.7%-26.3%) overlap than expected under random sampling (1% overlap), and diversity-based strategies’ overlap with other strategies was much closer to the baseline expectation (0.6%-2.6% overlap) (**Fig. 4B**). These results were observed across model architectures (**Supplementary Fig. 6**), though ATTN in yeast somewhat deviated from this trend with lower overlaps, consistent with its low performance. Notably, many of these inter-strategy overlaps were not substantially lower than overlaps between round configurations in the same strategy (**Fig. 3C-D**), suggesting that different uncertainty-based selection strategies identify largely overlapping sets of informative sequences.

**Figure 4:**
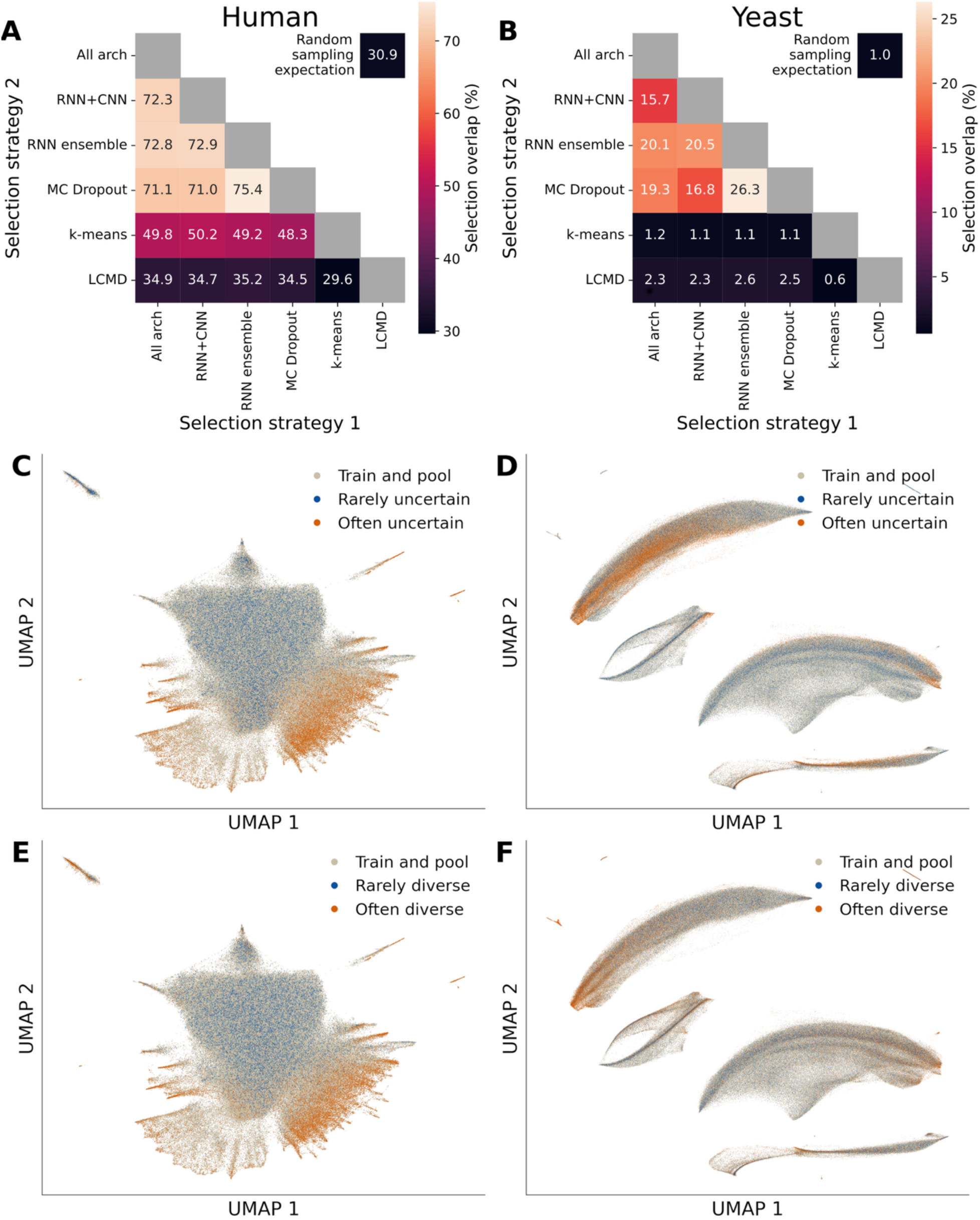
AL strategies achieve similar performance gains through partial overlaps in selection. **A-B)** Percentage of selected sequences shared (colors; numbers on heatmap; out of 60,000) between AL strategies (both axes), in human (**A**) and yeast (**B**) datasets. The expected overlap from random selection (**Methods**) is shown in the top-right corner. **C-F**) UMAP embeddings of the often (orange) and rarely (blue) selected sequences by uncertainty-based (**C-D**) and diversity-based (**E-F**) strategies, with the base training set and AL pool (gray), for the human (**C, E**) and yeast (**D, F**) datasets. All data shown are for RNN.

To better understand general trends in selection, we aggregated selections across strategies and model architectures for uncertainty and diversity-based strategies separately. In human, many sequences were selected by nearly all uncertainty-based strategies while a third of the pool was never selected (**Supplementary Fig. 7A**), indicating that models generally agreed on the uncertainty level of most sequences. Diversity-based selections also had more sequences frequently selected as compared to the expectation under random sampling, though to a lesser extent than uncertainty-based selections (**Supplementary Fig. 7B)**. In yeast, while most of the sequence pool was never selected (88.7% for uncertainty, 76.0% for diversity), many sequences were selected much more often compared to random sampling by both uncertainty and diversity-based strategies, indicating that a selection bias towards certain sequences persisted in a larger sequence space (**Supplementary Fig. 7C, D**).

Altogether, our analysis of the acquisition trajectories indicates that AL can consistently identify sequences that boost performance, selecting certain sequences substantially more frequently than random. Nevertheless, each comparison contains a substantial fraction of non-overlapping selections (24.6–70.4% in human; 73.7–99.4% in yeast), indicating that the strategies converge on comparable performance gains via somewhat distinct sample sets. This suggests that the space of informative sequences is large enough to support divergent acquisition trajectories. Indeed, very distinct selections can produce similarly strong results: k-means and LCMD both achieved similar performances that are superior to random sampling in yeast (**Fig. 2D, F, H**) despite having a sequence selection overlap that is less than expected by chance (**Fig. 4B**). This suggests that the similar performance gains of AL strategies can be achieved through diverse acquisition trajectories.

### AL selections are concentrated in specific regions of sequence space

We next wanted to examine where the sequences selected by AL sit in the sequence space. We embedded each sequence into a 2D UMAP projection by extracting the last layer embeddings of the sequence when passed through oracle models trained on the full datasets. We use the final-layer embeddings from each model to capture the regulatory properties driving expression since this layer is relatively low-dimensional (relative to previous layers) and necessarily includes the information needed to predict expression. Highlighting the sequences selected by each AL strategy, we observed that uncertainty- and diversity-based strategies selected from overlapping regions of sequence space in both datasets (**Supplementary Fig. 8**). We applied the same aggregation strategy used to define “often-uncertain” and “rarely-uncertain” sequences to diversity-based strategies, yielding “often-diverse” and “rarely-diverse” groups (see **Methods**).

We observed clear differences between the often and rarely selected sequences in both datasets. Often-selected sequences tended to be concentrated in smaller portions of the sequence space (**Fig. 4C-F**), while rarely selected sequences were substantially more scattered throughout the projection (**Fig. 4C-F**). This is consistent with our expectation: frequently selected sequences are more novel and lie further from the training distribution, while rarely selected ones tend to resemble many other sequences, sitting in denser, better-understood regions of sequence space. Interestingly, in both datasets, the often/rarely-uncertain sequences occupied similar sequence spaces as the often/rarely-diverse sequences (**Fig. 4C-F**). The overlap between often-uncertain and often-diverse sequences was 58.7% in human and 3.4% in yeast.

### AL methods preferentially select sequences with high transcriptional activity and certain regulatory features

Since we observed that often and rarely selected sequences occupy distinct regions of sequence space (**Fig. 4C-F**), we sought to characterize the selected sequences to understand what distinguishes these regions. Moreover, characterizing selected sequences also offers an opportunity to understand what features constitute informative training data, which could guide future experimental design and library construction. We systematically characterized their expression levels, ISM scores, TFBS content, and sequence composition.

We first examined the expression levels of selected sequences, since the AL pool sequences are already labelled using MPRAs. In both human and yeast, we observed that the often-uncertain and often-diverse sequences exhibited substantially higher expression levels than the rarely selected sequences, random selection, and the AL pool at large (**Fig. 5A-B**). We hypothesized that sequences with higher expression would be denser in expression-altering features encoded in the DNA sequence (e.g. TFBS). Thus, we performed ISM attribution analysis using models trained on the complete human and yeast datasets. In human, the often-selected sequences displayed significantly higher mean absolute ISM scores compared to the AL pool, while the rarely selected sequences displayed significantly lower mean absolute ISM scores (**Fig. 5C**), suggesting that selection was driven by a higher density of expression-altering sequence features. In yeast, this trend held only for uncertainty-based strategies, while the rarely/often-diverse sequences median ISM scores were similar to those of the AL pool and the random selection (**Fig. 5D**).

**Figure 5:**
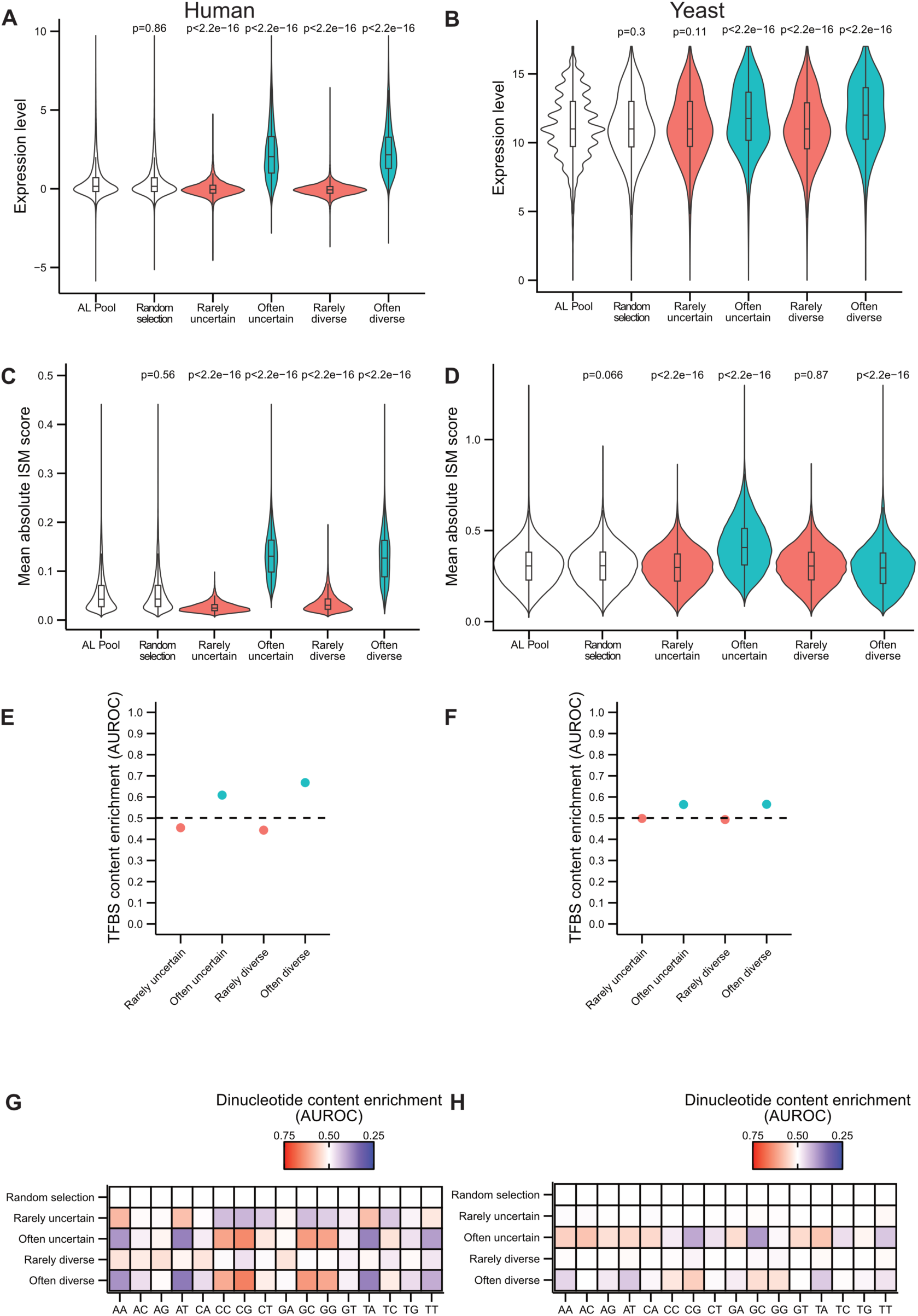
Characterization of rarely and often-selected sequences by AL methods. **A-B)** Expression levels (*y*-axis) of different sequence sets (*x*-axis) as measured by MPRAs of (A) human K562 sequences^51^ and (B) yeast sequences^52^ in the 3x20k configuration. P-values of the Mann-Whitney test between each sequence group and the AL pool are represented above the corresponding violin plot. **C-D)** Mean absolute ISM score (*y*-axis) of different sequence sets (*x*-axis) in the (C) human dataset and (D) yeast dataset in the 3x20k configuration. P-values of the Mann-Whitney test between each sequence group and the AL pool are represented above the corresponding violin plot. **E-F)** TFBS content enrichment *vs* the AL pool as scored with the Area Under the Receiver Operating Characteristic (AUROC, *y*-axis) for the rarely and often-uncertain/diverse (E) human sequences and yeast (F) in the 3x20k configuration (*x*-axis). Average AUROC for 15 randomly selected sequence sets compared to the AL pool is represented as a dotted line. **G-H)** Dinucleotide (*x*-axis) content enrichment (red tones) or depletion (blue tones) in the rarely and often-uncertain/diverse sequences (*y*-axis) relative to the AL pool as measured by the AUROC for (G) human and yeast (H) sequences in the 3x20k configuration.

To understand what sequence features might underlie these expression differences, we analyzed the TFBS content of the AL selected sequences. We scanned human and yeast sequences for predicted TFBSs using FIMO^54^ and compared their TFBS content to that of their original AL pool using the Area Under the Receiver Operating Characteristic (AUROC) as a non-parametric measure of the effect size. AUROC can be interpreted as the probability that a randomly drawn sequence from one set of selected sequences has a higher TFBS content than a randomly drawn sequence from remainder of the AL pool, where a value of 0.5 indicates no difference between groups and values approaching 0 or 1 indicate increasing separation. As expected, randomly sampling from the AL pool yields AUROC values around 0.5 in both human and yeast datasets, confirming that randomly selected sequences have the same TFBS content as the AL pool from which they are drawn (**Fig. 5E, F** and **Supplementary Note**). In both the human and yeast datasets, often-selected sequences exhibit AUROC values above 0.5 (**Fig. 5E, F** and **Supplementary Note**), indicating enrichment for TFBS content relative to the AL pool. For rarely-selected sequences, AUROC values fall below 0.5 in the human dataset but remain close to 0.5 in yeast (**Fig. 5E, F** and **Supplementary Note**), suggesting that AL preferentially excludes TFBS-depleted sequences in human.

To determine whether TFBS content increase in often-selected sequences was driven by specific TFs or reflected a broad increase across all of them, we performed TF enrichment analysis (**Methods**) comparing rarely and often-selected sequences against the AL pool. In both human and yeast, the TFBS enrichments appeared to be driven by specific TFs. In human, often-selected and often-diverse largely enriched for similar TFBSs, and these same TFBSs also tended to be depleted in rarely-selected sequences (**Supplementary Fig. 9A, C**). In yeast, overlapping sets of TFBSs were enriched in often-diverse and often-uncertain sequences, but often uncertain sequences also showed more depleted TFs than enriched TFs compared to the AL pool; rarely-selected sequences had minimal enrichment or depletion in yeast (**Supplementary Fig. 9B, D**). Altogether, those results indicate that the higher TFBS content in often-selected sequences compared to the AL pool is likely driven by a restricted set of TFs in both human and yeast. We also tested whether TF enrichment patterns reflected insufficient exposure during initial training by examining the TFBS frequencies in the training set of significantly enriched and depleted TFs. In general, what we found was consistent with this idea, with more enriched TFBSs in often-selected sequences being less common in the training set, and the opposite trend was seen for TFBSs that are enriched among rarely-selected sequences, but the trends were quite variable between yeast and human and between selection schemes (**Supplementary Fig. 10**). Another possibility is that the regulatory effects of certain TFs are intrinsically harder to learn, for example, due to more complex or context-dependent binding logic and thus sequences bearing them would be preferentially selected.

Finally, we examined the dinucleotide content of selected sequences. We computed the AUROC of randomly selected sequences and found, as expected, no meaningful differences to the AL pool in human and yeast datasets, whereas AL-selected sequences exhibited distinct compositional biases (**Fig. 5G-H**). In human, the often-uncertain and diverse sequences showed enrichment for GC-rich dinucleotides and depletion for AT-rich dinucleotides compared to the AL pool (**Fig. 5G** and **Supplementary Note**), in line with the higher GC content of strong human promoters relative to genomic background^55–57^. In yeast, the often-uncertain sequences showed GC content depletion and AT enrichment, while the often-diverse sequences showed the inverse pattern (**Fig. 5H** and **Supplementary Note**). The AT enrichment in uncertain sequences aligns with observations that yeast core promoters of strongly expressed genes tend to be AT-rich^58^.

Overall, sequences frequently selected by AL methods in the human dataset had higher expression levels, higher ISM scores, greater TFBS content, and biased nucleotide composition relative to the AL pool. The yeast dataset showed relatively attenuated selection bias trends, likely because the pool is composed of random sequences, which are generally more homogenous in their feature composition than genomic sequences. Finally, all the patterns observed in the 3x20k configuration held in the 1x60k configuration for both human and yeast sequences (**Supplementary Fig. 11** and **Supplementary Note**), indicating that the observed preferential selection for transcriptionally active sequences is a result of AL selection rather than a consequence of the iterative process.

### AL reliably differentiates informative sequences from less informative sequences for model improvements

Since we observed that AL strategies favor selecting certain sequences while avoiding selecting others (**Supplementary Fig. 7**), we investigated whether the most frequently selected sequences are also the most informative for model refinement. To test this, we trained RNN models using often and rarely selected sequences from uncertainty- and diversity-based strategies in the 1x60k configuration. To evaluate whether uncertainty-based and diversity-based selections could contribute complementary signal, we also evaluated a “combined” strategy containing equal numbers of frequently selected sequences from both approaches. In all cases, these selections were added to the initial model’s training set and compared against a randomly sampled set of the same size, and to the best individual uncertainty-based AL strategy (RNN ensemble, which had the highest average performance across datasets and test sets).

Models trained with the best-performing individual AL strategy achieved the highest performance across datasets and test sets, whereas aggregate approaches (often-uncertain, often-diverse, combined) sometimes matched but rarely exceeded this performance on all test sets (**Fig. 6A-B, Supplementary Fig. 13**). This result demonstrates that often-selected sequences are not inherently superior to the best individual strategy. One explanation is that individual strategies select sequences better tailored to their own model’s weaknesses compared to aggregate sets. Alternatively, they may be selecting more diverse (i.e. less redundant) sequence sets (**Supplementary Fig. 8**). The often-diverse sequences performed similarly to, though sometimes slightly weaker than, the often-uncertain sequences (**Fig. 6A-B**), consistent with the two groups selecting from similar regions of sequence space (**Fig. 4C-F**). This contrasts with the clearly better performance of individual uncertainty-based strategies over diversity-based strategies (**Fig. 2C-H**), and suggests that diversity-based strategies may benefit from running multiple times to identify recurrently selected sequences.

**Figure 6:**
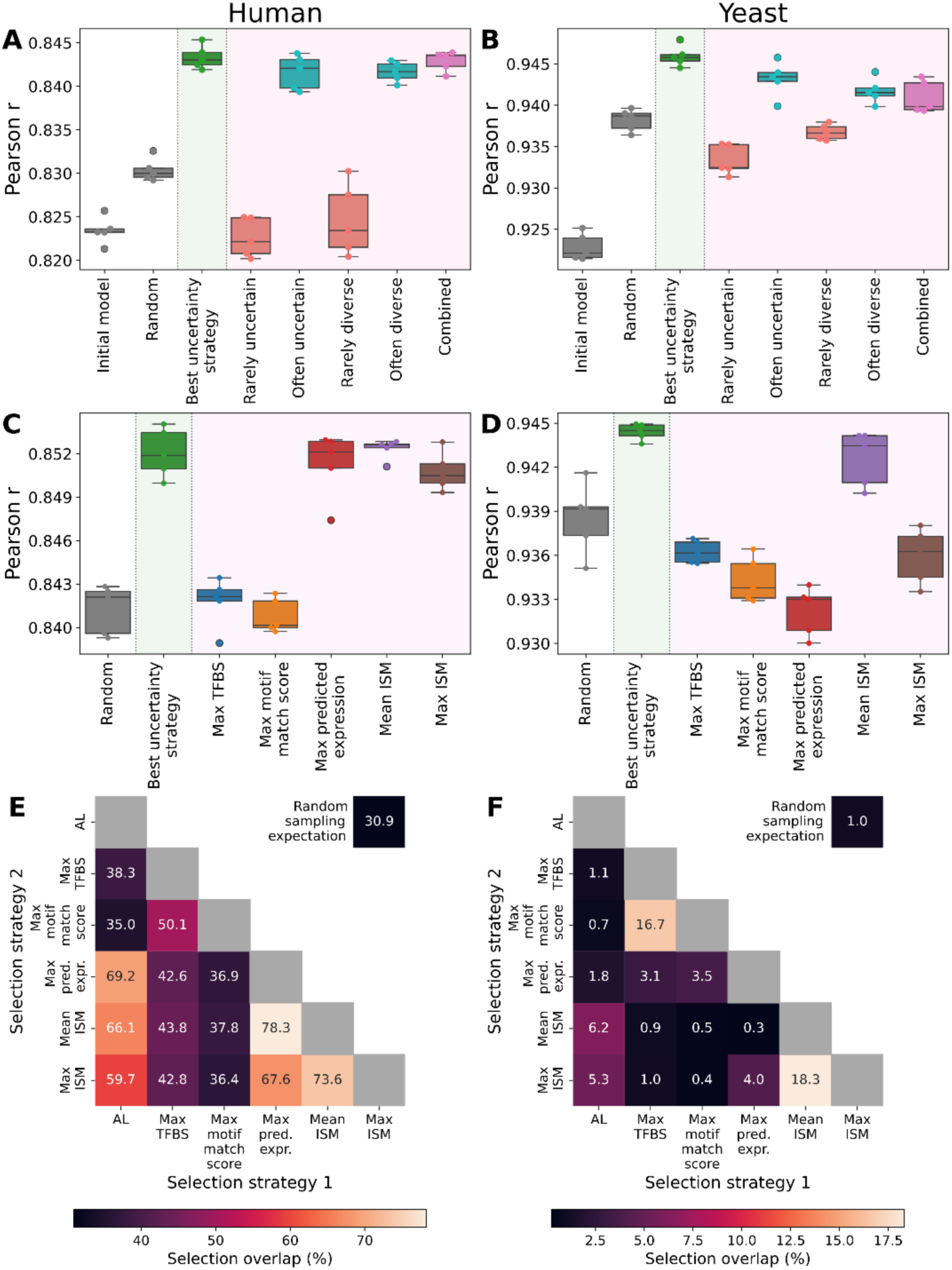
Sequences selected by the best AL strategy can outperform ensembles of different AL methods and biological acquisition functions. **A-B)** Performance (Pearson *r*, *y*-axis; ID test set), of models trained on the rarely- and often-selected sequences by AL (*x*-axis) in the human (**A**) and yeast (**B**) datasets, for uncertainty- and diversity-based strategies, and a “combined” strategy of half most uncertain and half most diverse sequences (pink background). Results are compared to the initial model and random selection (blank background) and to the strongest individual uncertainty-based strategy (green background). **C-D)** Performance (Pearson *r*, *y*-axes) on the ID test set for the best uncertainty-based strategy (green background) compared to random sampling (blank background) and five biologically driven sequence acquisition strategies (purple background; *x*-axis), for human (**C**) and yeast (**D**) datasets. All data shown are for RNN. **E-F)** Percentage of selected sequences (numbers on heatmap) shared between acquisition functions (both axes), in human (**E**) and yeast (**F**) datasets. The expected overlap from random selection (**Methods**) is shown in the top-right corner.

Models trained on rarely-uncertain and rarely-diverse sequences consistently underperformed the random baseline. In the human dataset, these models performed similarly to the initial model (**Fig. 6A**). In the yeast dataset, the much larger sequence pool meant most sequences were never selected (**Supplementary Fig. 7C-D**), and thus this selection method largely resembled random selection (**Fig. 6B**). These trends extended to VEP and OOD test sets (**Supplementary Fig. 13**), suggesting that rarely selected sequences contribute little additional information. Taken together, the poor performance of rarely selected sequences and the comparable performance of often-selected sequences relative to the best individual AL strategy (**Fig. 6A-B**) suggest that AL can efficiently distinguish informative from uninformative sequences.

We next asked whether often-selected sequences would carry enough signal to improve models even without the base training set. To test this, we trained models using the often-uncertain and often-diverse sequences as standalone training sets. In both datasets, the often-uncertain sequences underperformed random sampling about half the time, while the often-diverse sequences matched or outperformed the random baseline and the model trained with often-uncertain sequences on all tasks (**Supplementary Fig. 14**). These results indicate that often-uncertain sequences encode regulatory patterns underrepresented in the base training set, making them valuable as complementary data, but are not on their own a representative sample of the training distribution. By contrast, the stronger standalone performance of often-diverse sequences suggests that diversity-based selection captures a more representative subset. This distinction may partly explain the performance gap between the two strategy classes: uncertainty-based selection targets sequences the model finds hardest to predict, yielding large gains when combined with a strong initial model but performing poorly without one, whereas diversity-based selection produces a more balanced training set that is less dependent on the initial model but less targeted in what it adds.

### AL is more reliable than biologically-driven acquisition functions

Motivated by our observation that AL preferentially selected sequences with the features of high transcriptional activity (elevated expression levels, higher ISM scores, and high TFBS content) (**Fig. 5A-H**), we next asked whether directly selecting for these sequence characteristics could achieve comparable performance to AL. We tested five simple and intuitive biologically-driven approaches for selecting sequences from the selection pool: (1) highest TFBS content (how many TF binding sites a sequence contains); (2) maximum motif match score (how strongly a sequence matches each TF motif); (3) predicted maximum expression (using the initial model); and the highest (4) mean and (5) maximum absolute ISM scores (using the initial model; **Methods**). Each was used to select the top 60,000 sequences from the AL pool for model refinement. We compared these approaches to single-round uncertainty-based AL and to random selection.

Uncertainty-based AL consistently produced the largest gains across both datasets and tasks. In the human dataset, maximum predicted expression, mean and maximum absolute ISM selection approached the performance of the best uncertainty-based strategy and exceeded the random baseline, while both TFBS-based strategies (highest TFBS content and maximum motif match score) performed at or slightly below the random baseline (**Fig. 6C**). VEP and OOD results were similar (**Supplementary Fig. 15A, C**). In yeast, only the best uncertainty-based strategy and mean absolute ISM selection clearly outperformed random sampling, while the remaining biologically-driven strategies performed below the random baseline (**Fig. 6D**). While mean absolute ISM matched AL performance in the ID and OOD test sets, AL was clearly the strongest in VEP (**Supplementary Fig. 15B, D**). Taken together, these results suggest that model-derived uncertainty captures aspects of sequence informativeness that no single biological feature approximates reliably.

To understand what drove the performance differences among biologically-driven strategies, we examined the overlap between their selections and those of uncertainty-based AL (**Fig. 6E-F**). In human, ISM-based and maximum predicted expression strategies shared substantially more sequences with AL (59.7%–69.2%) than expected by chance (30.9%), while TFBS-based strategies overlapped with AL at rates only modestly above the baseline expectation (35.0%–38.3%) (**Fig. 6E**). In yeast, the same pattern held but at much lower absolute overlap given the larger pool: ISM-based strategies shared 5.3%–6.2% of sequences with AL, well above the 1.0% expected by chance, whereas TFBS-based strategies overlapped at near-chance levels (0.7%–1.1%) (**Fig. 6F**). Notably, the two TFBS-based strategies overlapped substantially with each other (50.1% in human, 16.7% in yeast) but shared little with AL or with the other biologically-driven approaches, indicating that they select a distinct and largely uninformative subset of sequences. This gradient in overlap tracked with performance: strategies that selected more similarly to AL also produced larger gains (**Fig. 6C-D**).

Our results indicate that while biological features such as TFBS content, expression levels, and ISM scores correlate with frequent AL selection, no single feature is sufficient to replicate the performance gains that uncertainty-based AL achieves. This suggests that sequence informativeness arises from a combination of properties that model-derived uncertainty captures implicitly but that simple biological heuristics do not.

## Discussion

Our study establishes AL as a robust strategy to improve sequence-to-expression models in lab-in-the-loop settings. Across diverse datasets, model architectures, and experimental configurations, uncertainty-based AL consistently outperformed random sampling, while diversity-based strategies yielded more variable results (**Fig. 2C-H, Supplementary Fig. 1C-H, 2C-H**). Notably, AL yields the largest performance improvements on generalization tasks, suggesting that it guides models toward learning general cis-regulatory logic rather than dataset-specific patterns (**Fig. 2E-H, Supplementary Fig. 1E-H, 2E-H**). This also suggests that evaluating AL on ID test data alone does not fully capture its utility, and test sets measuring generalization should be used to ensure a holistic evaluation. To support broad adoption of these methods, our accompanying framework, nextFrag, provides end-to-end tools for both AL selection and biological characterization of the selected sequences.

Based on these results, we outline practical guidelines for implementing AL in sequence-to-expression modeling.

First, uncertainty-based strategies emerged as the best performing approach, consistently outperforming diversity-based strategies across our benchmarks (**Fig. 2C-H, Supplementary Fig. 1C-H, 2C-H**). Performance was comparable across individual uncertainty-based strategies, including ensembles of varying size and architectural composition, and MC dropout, suggesting that the specific choice of uncertainty estimator is not critical within the configurations we tested. Additionally, uncertainty-based strategies are relatively computationally inexpensive, requiring only a small number of forward passes per sequence, whereas diversity-based methods such as k-means and LCMD require embedding all pool sequences and performing computationally intensive clustering. This efficiency gap becomes increasingly important as pool sizes scale to hundreds of millions of sequences, as anticipated in future experimental applications. Given the consistent performance advantage and favorable scaling properties, we recommend uncertainty-based strategies as the default choice for AL in this domain.

Second, we found that well-trained initial models are essential for effective AL. ATTN in yeast did not substantially outperform the random baseline, selecting less informative sequences than RNN or CNN **(Supplementary Fig. 2D, F, H)**. This observation suggests that starting AL from weak models, whether due to insufficient training data, poor architectural fit, or both, may be inefficient^59^. AL appears better suited to improving models that have already achieved reasonable baseline performance, after which it can efficiently guide further dataset expansion^59^. This dependence is particularly relevant for uncertainty-based strategies that select sequences away from the initial training distribution (**Supplementary Fig. 8**), containing less common sequence characteristics such as infrequently appearing TFs (**Supplementary Fig. 10**).

Third, a single, but larger AL round can substitute for multiple smaller rounds, with minimal performance loss **(Fig. 3A-B, Supplementary Fig. 4A-D)**. This is particularly relevant for lab-in-the-loop applications, where each AL round requires a full experimental cycle of sequence synthesis, MPRA measurement, and model retraining. Because experimental time increases linearly with each sequential AL round, whereas sequence synthesis costs increase sub-linearly for larger libraries, prioritizing fewer, larger experiments can substantially accelerate the experimental timeline and reduce costs while preserving the benefits of AL.

Beyond these practical recommendations, our results also offer insight into why AL works and what makes sequences informative for model training.

We visualized the most uncertain AL selected sequences, finding that they were often concentrated on the periphery of sequence space (**Fig. 4C-F**), thus providing complementary information instead of repeating examples similar to those in the training data (**Supplementary Fig. 10**). This finding is supported by our observation that models trained exclusively on often-uncertain sequences underperformed random sampling (**Supplementary Fig. 14**), indicating that selected sequences are only informative where an initial training set is also present.

We examined selection trajectories and found that the similar performance gains were driven by notable, but not complete, overlaps in selection (**Fig. 4A-B, Supplementary Fig. 6**). Moreover, models trained on often-uncertain and often-diverse sequences performed slightly worse than the model trained on sequences selected by the best performing AL strategy (**Fig. 6A-B**). Altogether, this indicates that there are several viable paths to improving model performance, with no single optimal set. Importantly, we also found that some sequences were avoided by AL selection (**Supplementary Fig. 7**), with these sequences tending to lead to minimal improvement (**Fig. 6A-B, Supplementary Fig. 13**). This shows that the sequence space contains regions of high redundancy that AL avoids.

To understand what biological features made for informative sequences, we characterized them and found consistent enrichment for biologically relevant features in both human and yeast. In human, AL favored sequences with higher GC content, elevated TFBS density, and higher expression levels, consistent with known properties of strong human promoters^55–57^ (**Fig. 5**). In yeast, uncertainty-based strategies selected sequences with higher expression levels, greater TFBS content, and AT-rich composition, aligning with characteristics of strong yeast promoters^58^ (**Fig. 5**). All the aforementioned patterns were consistent across the 3x20k and 1x60k configurations, confirming that AL preferentially selects for biologically relevant sequences regardless of round configuration **(Supplementary Fig. 11)**.

From the above observations, a natural question arises whether directly selecting for these biological features could bypass the need for model-guided AL altogether. Our results suggest not: while higher expression levels, high ISM scores, and TFBS density correlate with AL selection (**Fig. 5A-F**), directly optimizing for these features did not outperform AL (**Fig. 6C-D**). Strategies selecting sequences with maximal predicted expression or TFBS content performed inconsistently, occasionally matching AL but sometimes underperforming the random baseline, particularly on generalization tasks (**Fig. 6C-D, Supplementary Fig. 15**). Model-derived informativeness therefore captures aspects of sequence utility that extend beyond any single biological feature, reinforcing the value of AL as an integrated selection framework for lab-in-the-loop settings.

We acknowledge several limitations that constrain the generalizability of our findings and may require further exploration. First, in our benchmark we simulated AL by subsampling sequences from individual large datasets, which cannot capture the experimental factors such as batch effects and measurement noise that may differ between AL rounds. Second, we focused on short regulatory sequences (80-200 bp); whether these findings generalize to genome-scale models^1,2,8,9,16,17,60–62^ remains to be explored. Third, our selection pool was limited to sequences that were already present within each dataset. Since any DNA sequence can theoretically be synthesized and tested, future work should incorporate diverse sequence generation approaches^51,63–70^, including targeted mutagenesis, motif-based design, and generative models to construct larger and more diverse pools that allow exploration of sequence space beyond the initial labeled data. Such expanded pools, particularly those constructed by generative models, may reveal different selection dynamics across strategies and enable fully generative lab-in-the-loop framework where pool generation and sequence selection are jointly optimized.

## Methods

### Experimental Procedure

#### Datasets

We performed our benchmark on the Random Promoter DREAM challenge dataset^52^ (“yeast”), containing 6,739,249 80-bp random sequences, and the Gosai et al. K562 MPRA dataset^51^ (“human”), containing 367,364 200-bp reference allele genomic sequences. These datasets were chosen for their large sizes and different nature of training data (genomic and random, respectively), as well as the availability of test sets spanning multiple distributions, to better measure performance across sequence space and measure overfitting to training data. Both datasets provide a scalar expression value for each sequence. In human, we used the expression value measured in K562 only (K562_log2FC).

#### Preprocessing

For yeast, each 80 bp random sequence was padded to 150 bp with its corresponding plasmid (57 bp on the 5’ end and 13 bp on the 3’ end), as done in the DREAM challenge models^52^. For human, sequences shorter than 200 bp were padded equally on both ends with Ns. Sequences were one-hot encoded, with Ns encoded as the absence of a nucleotide (i.e. a zero vector). Reverse complements were added to each dataset, with an additional channel specifying whether a sequence was in the forward or reverse orientation. At inference time, models predicted on both forward and reverse strands and the predictions were averaged. For yeast specifically, a sixth channel was added to represent whether the expression was a singleton (i.e. integer value), with this channel set to 0 (i.e. non-singleton) at inference time. Model inputs had shape (batch_size,150,6) for yeast and (batch_size, 200, 5) for human.

#### Data splits

In yeast, our train/validation/test split matches the Rafi et al. paper. A randomly chosen 20,000 sequence subset of the validation split was used as the validation set, given the heavily down sampled training data. Our final split contained 100,000 training, 5,965,324 pool, 20,000 validation, and 71,103 test sequences. In human, we used hashFrag^18^ to generate orthogonal train/validation/test splits (ratio 80:10:10) to minimize homology-based data leakage. Our final split contained 100,000 training, 193,890 pool, 36,737 validation, and 36,737 test sequences.

#### Test sets

Models were evaluated on three test sets: ID, VEP, and OOD, with ID and OOD measuring ability to predict expression and VEP measuring ability to predict difference in expression. ID test sets were the DREAM Challenge “random” test subset for yeast, and the hashFrag test split for human; these sets are designed to measure model performance within the distribution of the training data. VEP test sets were the DREAM Challenge “SNV” test subset for yeast (44,340 pairs), and the difference between reference and alternate alleles in the hashFrag-split test set for human, where available (34,337 pairs). This measured the models’ ability to predict difference in expression, in contrast to directly predicting expression as in the other test sets. OOD test sets were the DREAM Challenge “native” test subset for yeast (997 sequences), and sequences designed for high K562-specific expression in human (sequences labeled with target_cell=k562 and origin= ‘AdaLead’ or ‘FastSeqProp’ or ‘Simulated_Annealing’ in the cell-type-specific CRE library^51^; 16,892 sequences), designed to measure model understanding of cis-regulatory logic more generally.

#### Models

We used the DREAM Challenge model architectures (CNN, RNN, ATTN)^52^. We made the following modifications to allow using MC dropout as a sequence selection strategy: CNN had dropout added to its core block, and ATTN had dropout added to its first layers block. Dropout probability was set to 0.1. RNN already had dropout and was unmodified.

#### Training

Training was performed for 80 epochs with a batch size of 32 for the human dataset and 256 for the yeast dataset. We used the AdamW optimizer (weight decay 0.01, learning rate 0.005 for RNN and CNN, 0.001 for ATTN) and a one cycle LR scheduler with cosine annealing. Model performance was evaluated using Pearson r on the validation set, with the best performing model on this metric being selected.

#### Evaluation

For ID and OOD test sets, models were evaluated using the Pearson correlation between model predictions and the respective test set. For VEP, models predicted on both alleles and the Pearson correlation between the predicted and actual differences in expression was used.

### Description of Sequence Selection Strategies

#### Uncertainty-based

##### Ensemble

An ensemble of models was trained using the same training set. Each model ran inference on the unlabeled pool, and the variance between model predictions was calculated for each sequence. For *k* sequences selected per round, the *k* highest variance samples were selected. We tested ensembles differing in composition and size: 3 models total with one of each architecture (RNN, CNN, ATTN, referred to as “All arch”), 2 models total with one RNN and one CNN as these were our two best performing architectures, and 5 models of the same architecture (e.g. 5 of RNN referred to as “RNN ensemble”).

##### Monte Carlo (MC) dropout

Each model independently selected sequences according to MC dropout^41^. Briefly, this involved enabling dropout at inference time, then making 5 forward passes on each sequence and calculating the variance between the predictions for each sequence. The *k* highest variance samples were selected for labeling.

#### Diversity-based

##### Abstract sequence embedding

for both diversity-based strategies, we employed the last layer embedding technique common in deep AL to create an abstract embedding of our DNA sequences^44,45,71,72^. This allows the creation of a notion of sequence similarity, critical for diversity-based methods. Models ran inference on all pool sequences and the output before the final mapper layer was extracted and concatenated into a single dimension. To reduce computational cost of clustering, principal component analysis with whitening was performed with the top 64 components selected and used for diversity-based strategies.

##### k-means++

k-means clustering was performed using the k-means++ initialization^73^, with *k* clusters (as many clusters as the number of selected sequences). For each cluster, the closest sequence to the cluster center (measured by Euclidean distance) was selected for labeling.

##### Largest Cluster Maximum Distance (LCMD)

we used the LCMD selection algorithm described in Holzmüller et al.^44^, which prioritizes maximally different sequences from previously selected ones. Briefly, a random point is selected as the first cluster center, then the following algorithm is applied: the largest cluster (as measured by the sum of Euclidean distances from all elements within the cluster, to the cluster center) is identified, and the furthest point from the cluster center within that largest cluster is chosen as a new cluster center. The identity of the largest cluster is updated, and the process is repeated for *k* iterations, before all cluster centers are selected for labeling.

### Characterization of sequences selected by AL

#### Selection frequency between AL strategies

Each distinct selection strategy was counted: one for each architecture and replicate for MC dropout, k-means, and LCMD (15 per method); one for each replicate for “all arch” and RNN+CNN (5 per method), one for each architecture for the single architecture ensemble since they will have the exact same selection across replicates (3 total). This gives 28 total uncertainty and 30 total diversity sampling selections. This was done separately in the 3x20k and 1x60k configurations. For each sequence, the number of replicates in each category (3x20k uncertainty, 3x20k diversity, 1x60k uncertainty, 1x60k diversity) that selected it in any round was counted.

#### Baseline sequence set overlap calculation

We calculate the proportion of sequences selected by both strategies, given that the sequence was already selected by the first strategy. For the human dataset, each strategy selected 60,000 sequences from a pool of 193,890 sequences, corresponding to 31% of the pool. The expected overlap between two independent selected sets is therefore 31%, as each sequence selected by the first strategy has a 31% probability of also being selected by the second. For the yeast dataset, strategies selected 60,000 sequences from a pool of 5,965,324 sequences, approximately 1% of the pool. By the same reasoning, the expected overlap under random selection is approximately 1%.

#### Selection of most and least uncertain/diverse sequences

Sequences were ranked by selection frequency across uncertainty- or diversity-based AL strategies. “Most uncertain/diverse” sequences were defined as those most frequently selected, whereas “least uncertain/diverse” sequences were defined as those least frequently selected. For the human dataset, the most and least uncertain/diverse subsets were defined as the top and bottom 10% of sequences ranked by selection frequency for each AL strategy, yielding 19,389 sequences per strategy per configuration. For the yeast dataset, a more stringent threshold was applied, selecting the top and bottom 1% of sequences by selection frequency, yielding 59,654 sequences per strategy per configuration. For the combined approach, the top half of the most uncertain/diverse sequence subsets were used. When overlap occurred, remaining sequences were selected based on highest combined selection frequency across uncertainty- and diversity-based strategies.

#### UMAP projections

Sequences were passed through an oracle RNN model trained on the base training set and full AL pool, and last layer embeddings were extracted. To reduce computational cost of UMAP, PCA with whitening was performed and the top 64 components were kept and input into UMAP.

#### In silico mutagenesis analysis

All possible single-nucleotide mutations were applied to each input sequence, yielding 600 mutations per sequence for human and 240 for yeast; in yeast, only the 80-bp random promoter region was mutated, with flanking sequence held constant. Oracles trained on the entire respective dataset ran inference on each mutated sequence, and the change in output relative to the unmutated sequence was computed for each mutation. Attribution scores were defined as the absolute change in output at each position. The mean absolute score was computed by averaging absolute attribution scores across positions, and the maximum absolute score was the largest absolute attribution for any position.

#### Dinucleotide content analysis

Dinucleotide frequencies for all human and yeast sequences were computed using the polygraph.sequence.kmer_frequencies() function from the Polygraph Python package^74^.

#### Transcription factor binding site scanning

TFBS scanning was performed using FIMO from the MEME suite^75^. For human sequences, position weight matrices (PWMs) in MEME format were obtained from the HOCOMOCO v13 database^76^ and filtered for TFs expressed in K562 cells (TPM > 1) using GTEx whole blood expression data^77^ (GTEx_Analysis_2017-06-05_v8_RNASeQCv1.1.9_gene_median_tpm). For yeast, PWMs were downloaded from JASPAR^78^ (Fungi non-redundant) and filtered for yeast TFs using the sacCer3 NCBI RefSeq annotation. Background models were generated with the fasta-get-markov tool from the MEME suite using the sequences from the human and yeast AL pools. For both species, only the best-scoring motif from either strand was retained using the --max-strand option, and motifs were called at FDR < 0.01. The total number of TFBS per sequence was computed with a custom R script.

#### Compositional difference scoring

To quantify compositional differences between selected sequences and the AL pool, we employed Cliff’s delta (Cd), a non-parametric effect size measure^79^, then we mapped Cliff’s delta back to the probability space by its linear relationship with AUROC. Dinucleotide frequency distributions in selected sequences were compared to the AL pool using the cliffs_delta() function from the effectsize R package (v1.0.1). For TFBS content, we compared the distribution of TFBS counts per sequence between selected sets and the AL pool. To discriminate between enrichment or depletion of the above sequence features, we performed 15 random selections of 20,000 and 60,000 sequences from the human and yeast AL pools respectively. We computed the Cd of each random set individually and averaged it to get the random selection for dinucleotide content or TFBS content. Finally, to put our effect size values into the probability space we transformed the Cd values into AUROC with the following formula: AUROC=(C_d_ + 1) / 2.

#### TFBS enrichment analysis

TFBS enrichment was assessed by computing Fisher’s exact test for each TF’s motifs counts obtained from the TFBS scanning with FIMO between the rarely/often-selected sequences and the AL pool. The test was deemed significant for FDR < 0.05 and enrichment was called for log2 odds ratio > 1 and depletion for log2 odds ratio < -1.

#### Motif match score

Each PWM was converted to a position-specific log-odds matrix by computing log₂ of the per-position nucleotide probability against a uniform background frequency of 0.25 per base. A reverse-complement counterpart was constructed for each motif to enable scoring on both strands. Each sequence was scanned against every motif at all valid sliding window positions, with the score at a given position computed as the sum of log-odds contributions across the motif width. The motif match score for a sequence–motif pair was defined as the maximum log-odds score across all positions and both strands, yielding a sequence × motif matrix of best-match scores used in downstream analyses.

#### Biologically driven acquisition functions

For maximum TFBS selection, refer to “Transcription factor binding site scanning,” with the 60,000 sequences with the highest TFBS count selected for addition to the training set. For maximum predicted expression selection, the initial model was used to predict expression on all pool sequences, and the top 60,000 sequences were selected. For ISM-based selection, the initial model was used to perform ISM and the 60,000 sequences with highest mean or maximum ISM scores were selected.

## Author Contributions

A.M.R. and C.G.D. conceived the study. A.M.R., J.Q., and E.C. designed experiments. J.Q. developed the codebase, trained models, and conducted the AL experiments. E.C. characterized the AL selected sequences. J.Q., A.M.R., and E.C. interpreted results. J.Q., E.C., and A.M.R. wrote the manuscript. C.G.D. edited the manuscript. C.G.D. and A.M.R. supervised the study. C.G.D. provided funding.

## Code Availability

Open-source code for nextFrag is available on GitHub at https://github.com/de-Boer-Lab/nextFrag and https://github.com/de-Boer-Lab/nextFrag-PaperCode.

## Data Availability

Data used for this study are available from Zenodo (https://doi.org/10.5281/zenodo.20094954). The human MPRA dataset is available from Zenodo (https://doi.org/10.5281/zenodo.10698014). The yeast MPRA dataset data is available from Zenodo (https://doi.org/10.5281/zenodo.10633252).

## Supplementary Figures

**Supplementary Figure 1:**
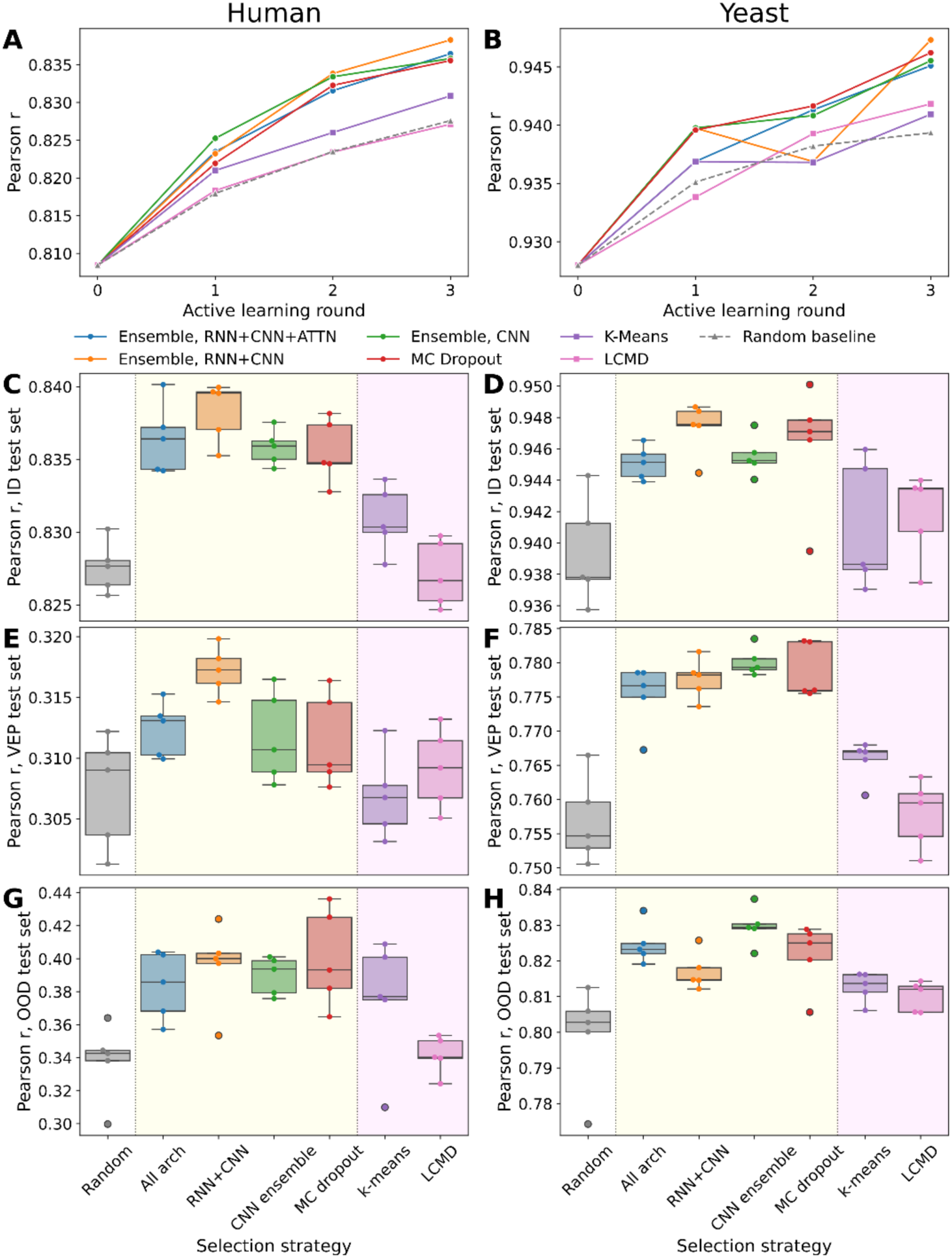
AL outperforms random sampling for CNN across test sets. **A-B)** Performance of CNN models (Pearson *r*, *y*-axes) across AL rounds (*x*-axes) for different selection strategies (solid lines) compared to a random sampling baseline (dashed gray line) on the ID test set, averaged across 5 replicates of CNN for human (**A**) and yeast (**B**) datasets. **C-H)** Performance of CNN models (Pearson *r*, *y*-axes) across AL selection methods (*x*-axes; yellow background for uncertainty- and magenta background for diversity-based strategies) compared to a random sampling baseline (blank background) on the ID test set **(C-D)**, VEP test set **(E-F)**, and OOD test set **(G-H)**, for human **(C, E, G)** and yeast **(D, F, H)** datasets.

**Supplementary Figure 2:**
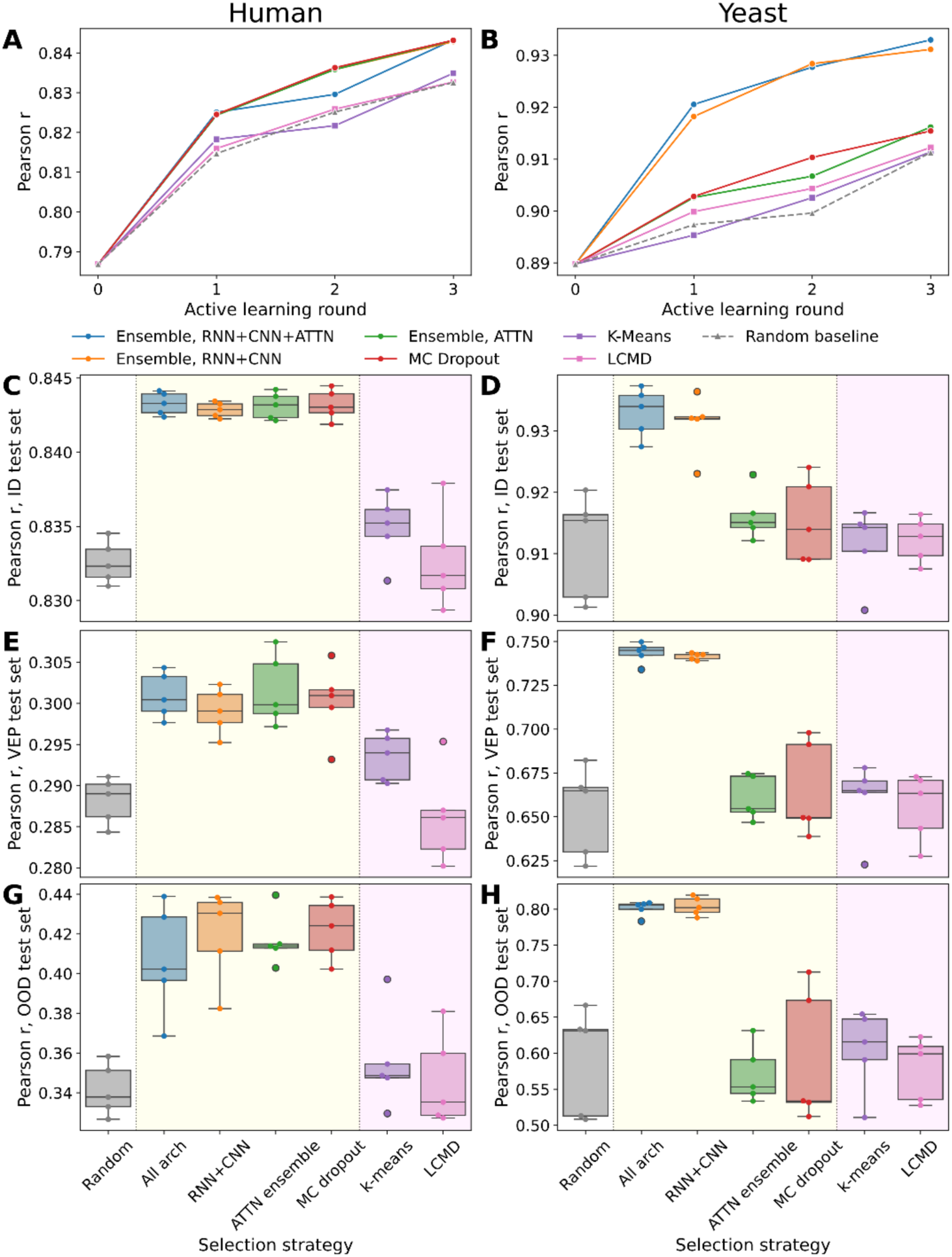
AL outperforms random sampling for ATTN across test sets. **A-B)** Performance of ATTN models (Pearson *r*, *y*-axes) across AL rounds (*x*-axes) for different selection strategies (solid lines) compared to a random sampling baseline (dashed gray line) on the ID test set, averaged across 5 replicates of ATTN for human (**A**) and yeast (**B**) datasets. **C-H)** Performance of ATTN models (Pearson *r*, *y*-axes) across AL selection methods (*x*-axes; yellow background for uncertainty- and magenta background for diversity-based strategies) compared to a random sampling baseline (blank background) on the ID test set **(C-D)**, VEP test set **(E-F)**, and OOD test set **(G-H)**, for human **(C, E, G)** and yeast **(D, F, H)** datasets.

**Supplementary Figure 3:**
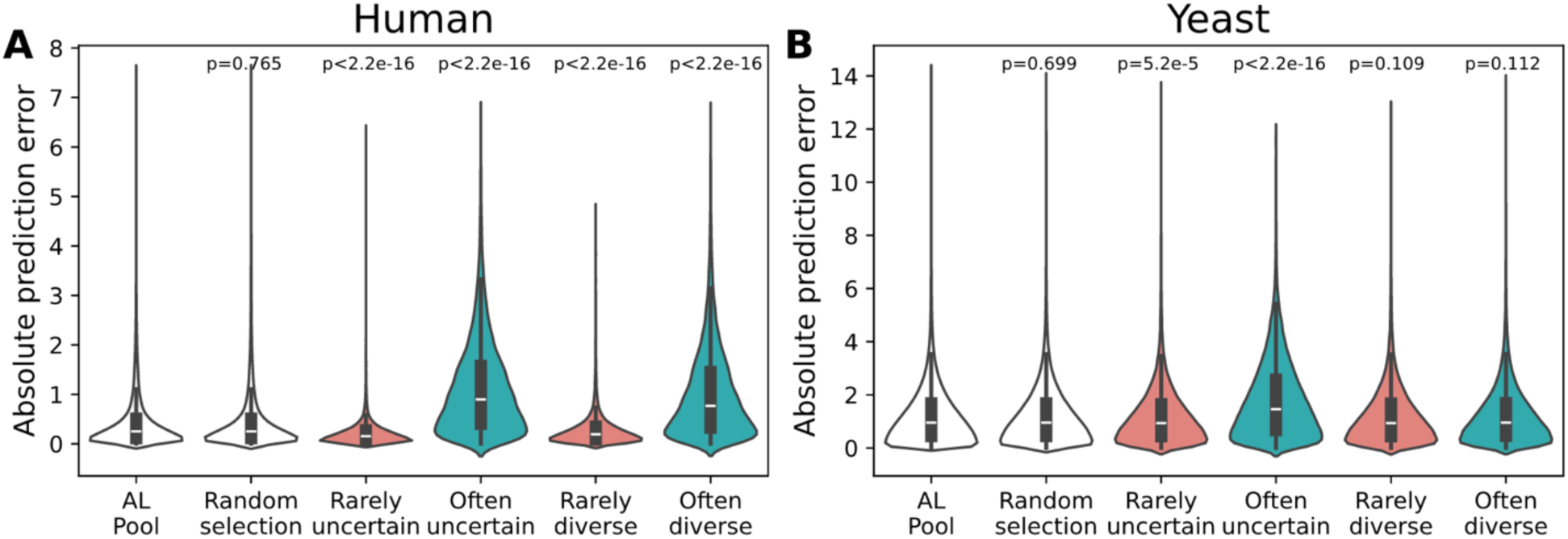
Prediction error for often and rarely-uncertain and diverse sequences. Absolute prediction error (*y*-axes) of different sequence sets (*x*-axes) in the human (**A**) and yeast (**B**) datasets. P-values of the Mann-Whitney test between each sequence group and the AL pool are shown above the corresponding violin plot.

**Supplementary Figure 4:**
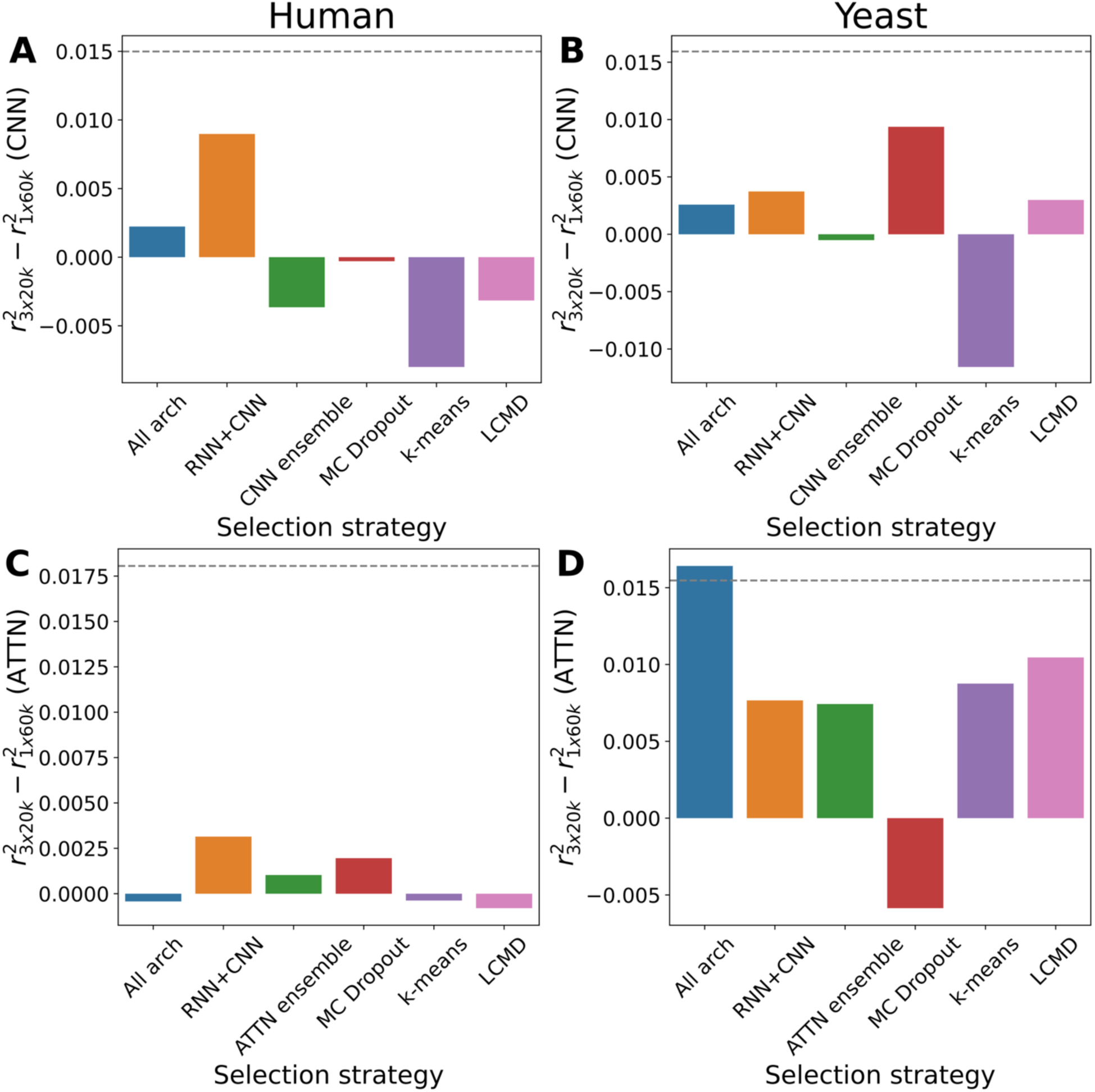
AL performance is usually similar between 3x20k and 1x60k configurations. Difference in median *r*^2^ between 3x20k and 1x60k configurations (*y*-axes), broken down by selection strategy (*x*-axes) for CNN (**A-B**) and ATTN (**C-D**) architectures, in the human (**A, C**) and yeast (**B, D**) datasets. Average difference from random sampling is indicated by the gray dashed line.

**Supplementary Figure 5:**
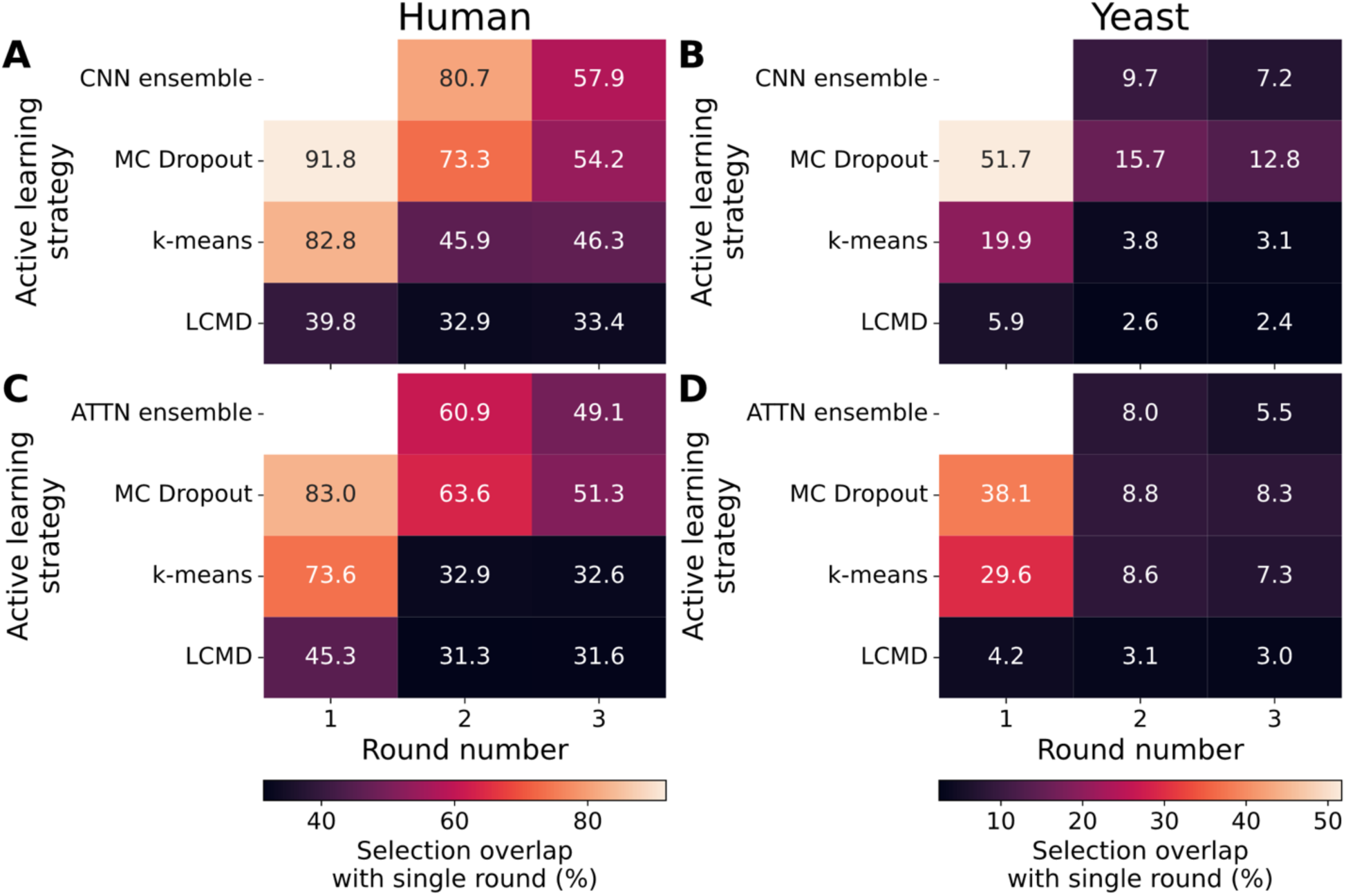
Selection similarity between 3x20k and 1x60k configurations across rounds. Proportion of sequences for each round (*x*-axes) of the 3x20k configuration that are shared with the corresponding 1x60k selection, for each selection strategy (*y*-axes), for CNN (**A-B**) and ATTN (**C-D**) architectures and human (**A, C**) and yeast (**B, D**) datasets. Round 1 for ensemble methods (CNN/ATTN ensemble) is not shown since selection is deterministic in that round.

**Supplementary Figure 6:**
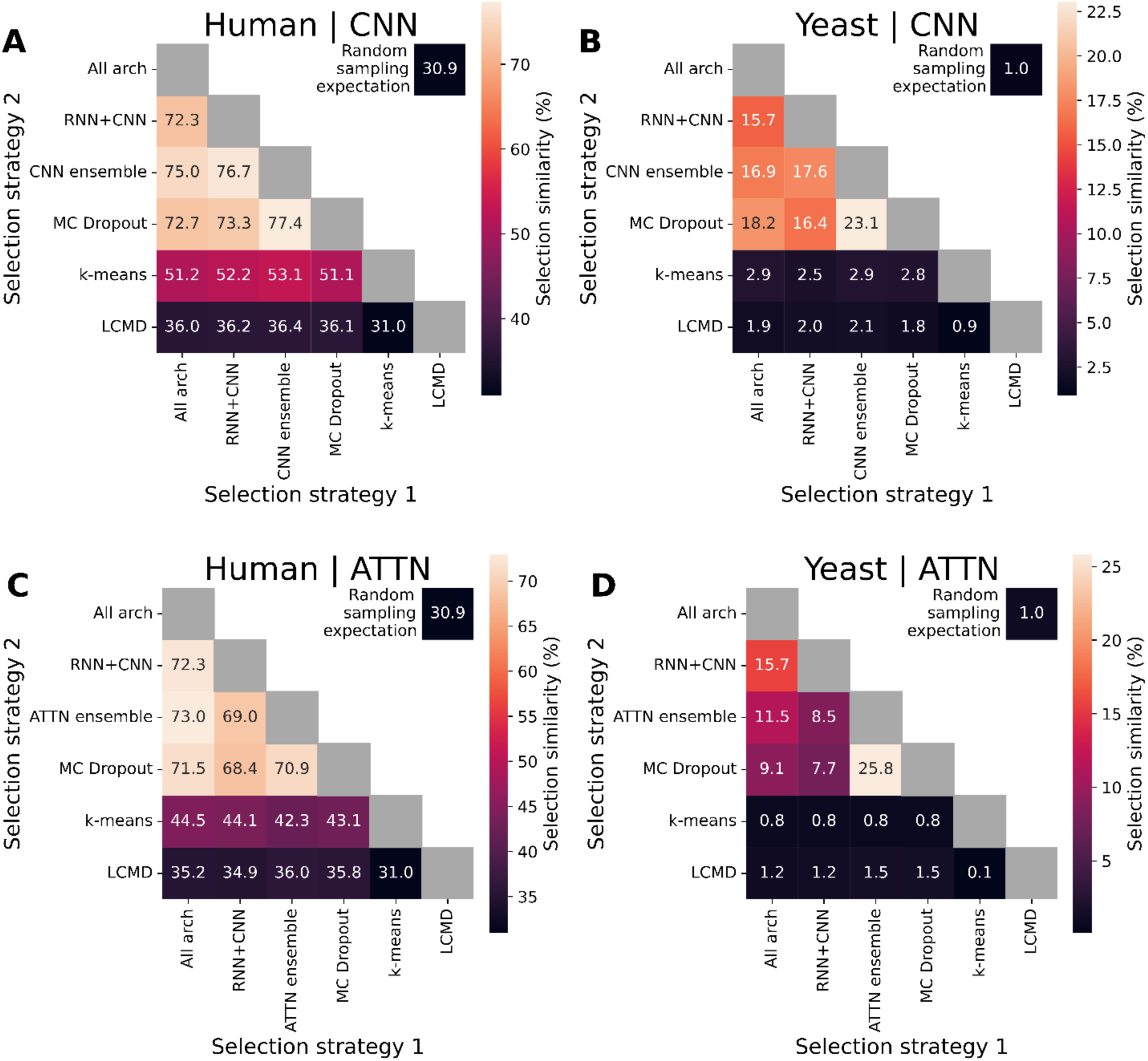
Comparison of AL strategies’ selections across different model architectures. Percentage of selected sequences shared (colors; numbers on heatmap; out of 60,000) between AL strategies (both axes), for CNN (**A-B**) and ATTN (**C-D**) architectures, in human (**A, C**) and yeast (**B, D**) datasets. The expected overlap from random selection (**Methods**) is shown in the top right corner.

**Supplementary Figure 7:**
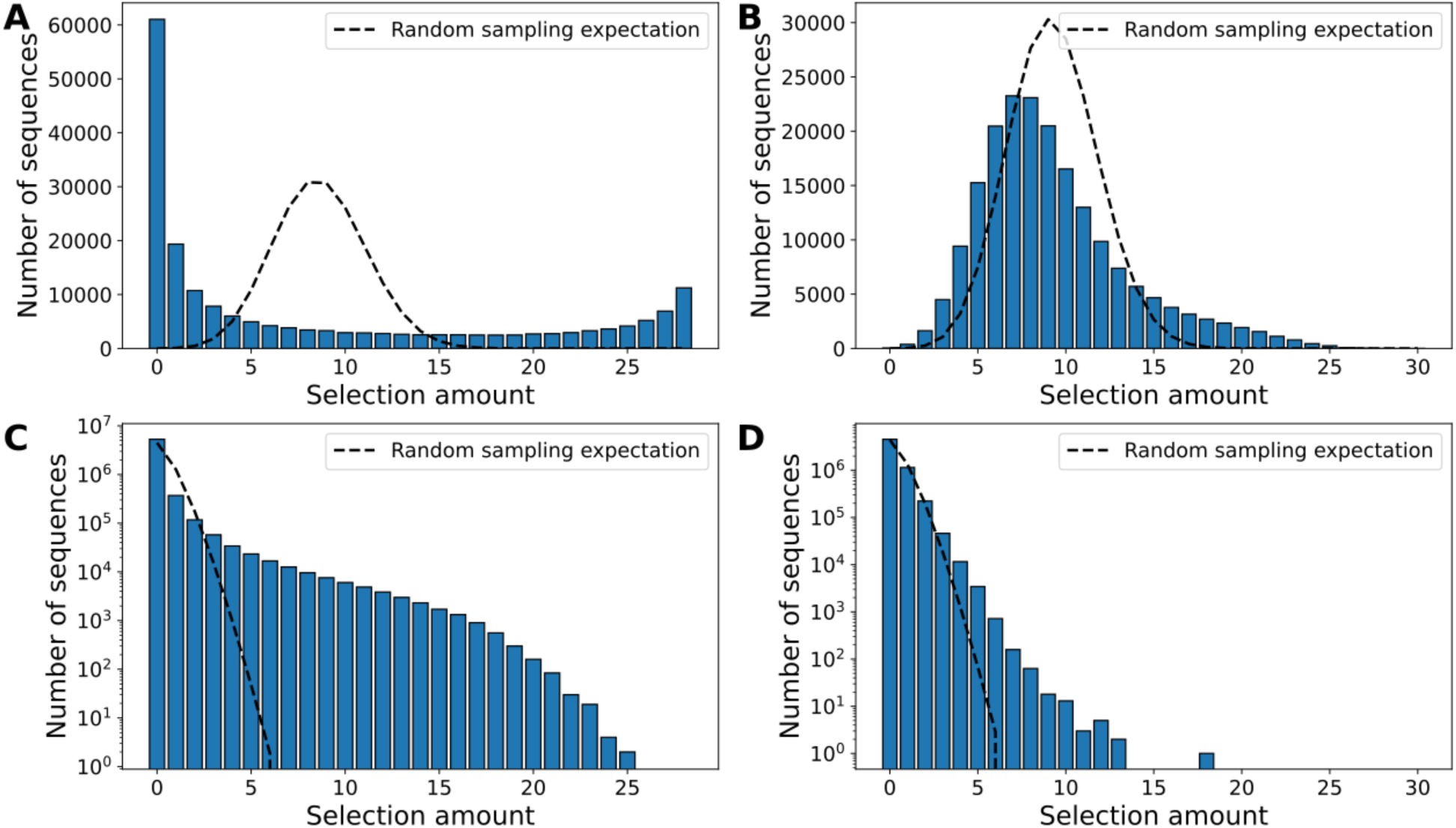
Selection frequency histograms across all AL strategies. Histograms of selection frequency (*x*-axes) across strategies and replicates in the human (**A-B**) and yeast (**C-D**) datasets, for uncertainty-based (**A, C**) and diversity-based (**B, D**) methods. The expectation from random sampling (binomial distribution), is shown as a black dashed line. Note the log scale on the *y*-axis in (**C, D**), due to large differences in order of magnitude. A bar at 60,000 for 0 selection amount means that 60,000 sequences were not selected by any replicate.

**Supplementary Figure 8:**
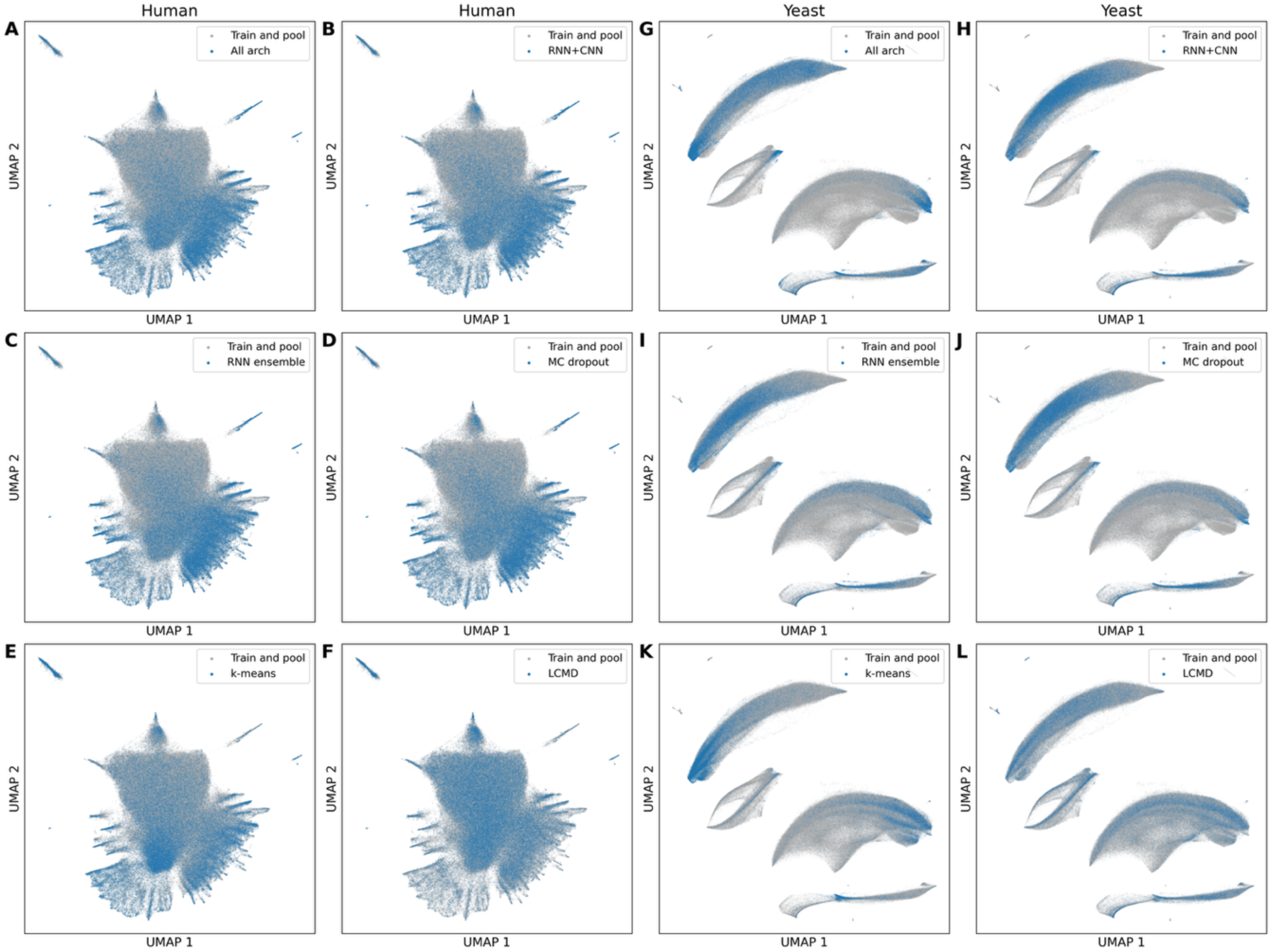
AL strategies selection patterns across the sequence space. UMAP embeddings of sequences selected by each AL strategy (blue), plotted against the base training set and AL pool (gray), for the human (**A-F**) and yeast (**G-L**) datasets.

**Supplementary Figure 9:**
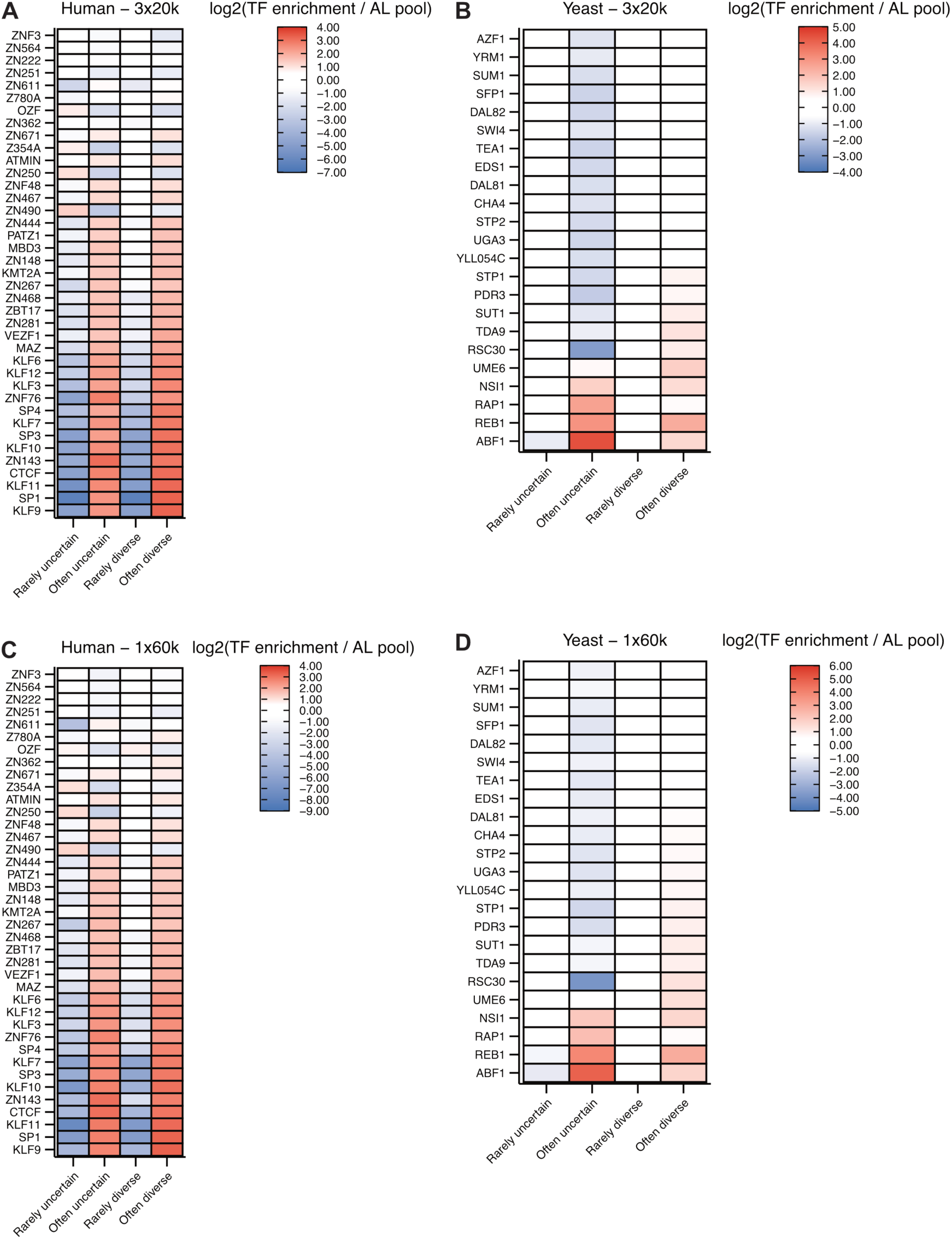
Per-transcription factor enrichment in selected sequence sets over the AL pool. **(A-D)** Enrichment (red tones) or depletion (blue tones) of TFs (*y*-axis) in often and rarely selected sequences by uncertainty and diversity methods (*x*-axis) relative to the AL pool sequences in the human dataset in the 3x20k (A) and 1x60k (C) configurations and in the yeast dataset in the 3x20k (B) and 1x60k (D) configurations. Enrichment or depletion is scored by the log2 odds ratio of the Fisher’s exact test between the occurrences of a given TF in selected sequences *vs* the AL pool. Only TFs with counts significantly different from the AL pool (FDR < 0.05) in at least one sequence set are represented.

**Supplementary Figure 10:**
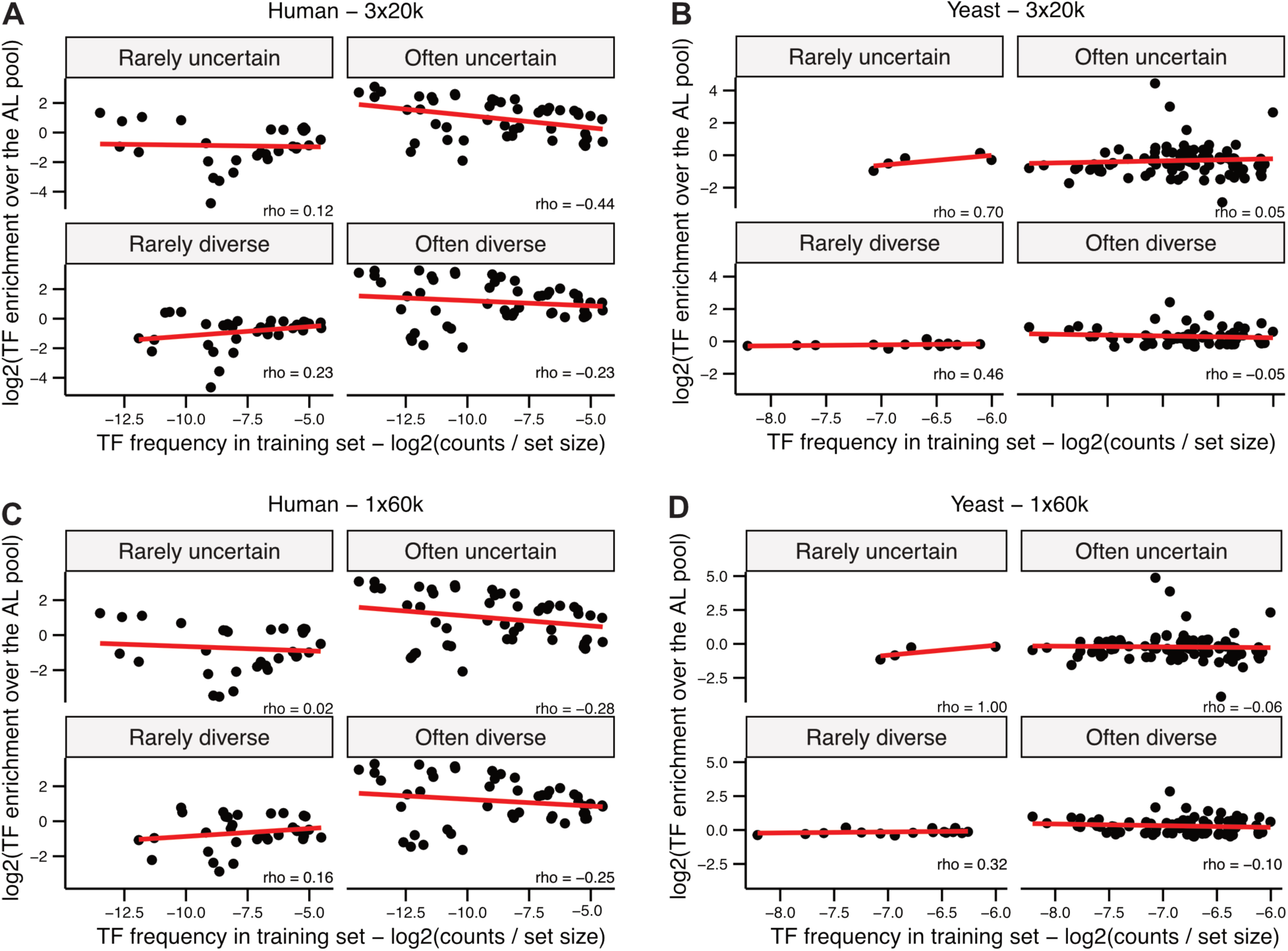
Frequency in the training set of significantly enriched TFs in selected sequences compared to the AL pool. **A-D)** TFs frequencies (*x-*axis) in training set sequences and their enrichment (log2 odds ratio from the Fisher’s exact test for individual TFs counts between the selected sequence sets and the AL pool) in often and rarely selected sequences by uncertainty and diversity AL methods relative to the AL pool (*y-*axis) for the human dataset in the 3x20k (A) and 1x60k (C) configurations and for the yeast dataset in the 3x20k (B) and 1x60k (D) configurations. Only the TFs with a Fisher’s exact test FDR < 0.05 are plotted. Red lines indicate linear fits. The coefficients (rho) for the Spearman correlation between the TF frequencies distribution and the OR distribution for each sequence sets are shown in each facet.

**Supplementary Figure 11:**
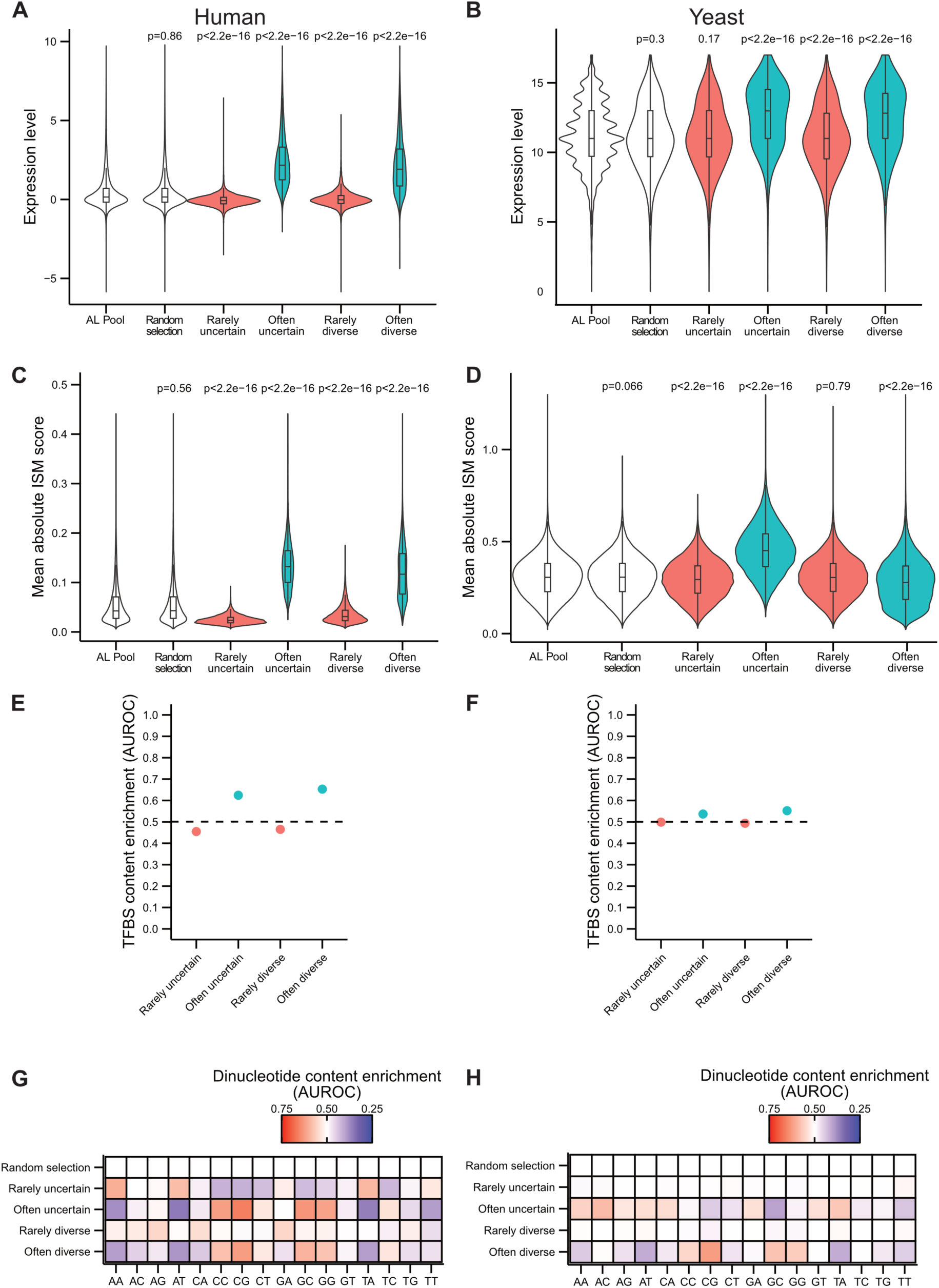
Characterization of rarely and often-selected sequences by AL methods in the 1x60k configuration. **A-B)** Expression (*y*-axis) of different sequence sets (*x*-axis) of (A) human K562 sequences^51^ and (B) yeast sequences^52^ in the 1x60k configuration. P-values of the Mann-Whitney test between each sequence group and the AL pool are represented above the corresponding violin plot. **C-D)** Mean absolute ISM score (*y*-axis) of different sequence sets (*x*-axis) in the (C) human dataset and (D) yeast dataset in the 1x60k configuration. P-values of the Mann-Whitney test between each sequence group and the AL pool are represented above the corresponding violin plot. **E-F)** TFBS content enrichment *vs* the AL pool as scored with the Area Under the Receiver Operating Characteristic (AUROC, *y*-axis) for the rarely and often-uncertain/diverse (E) human sequences and yeast sequences (F) in the 1x60k configuration (*x*-axis). Average AUROC for 15 randomly selected sequence sets compared to the AL pool is represented as a dotted line. **G-H)** Dinucleotide (*x*-axis) content enrichment (red tones) or depletion (blue tones) in the the rarely and often-uncertain/diverse sequences (*y*-axis) relative to the AL pool as measured by the AUROC for (G) human and yeast (H) sequences in the 1x60k configuration.

**Supplementary Figure 12:**
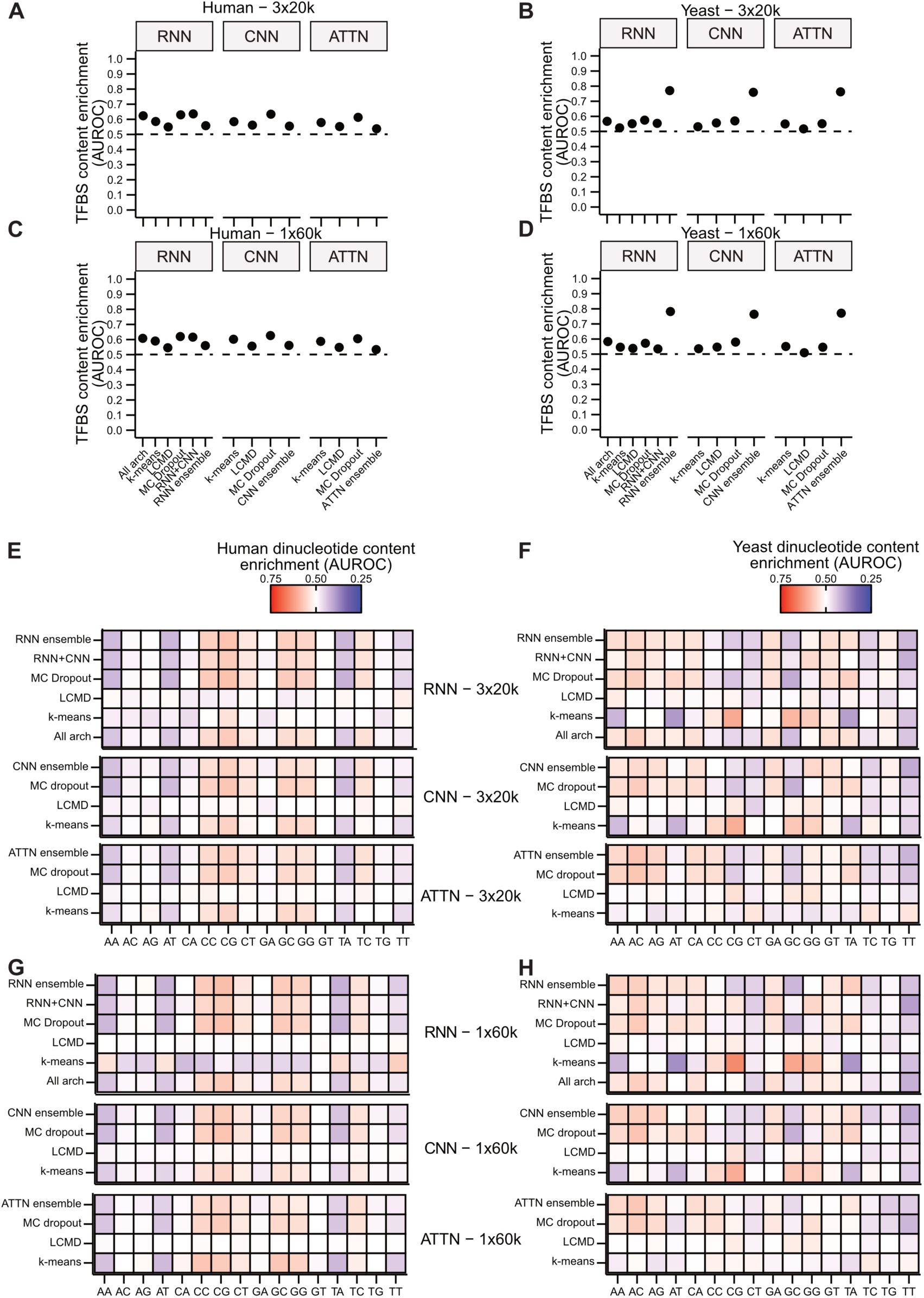
TFBS content bias and dinucleotide content bias of selected sequences across model architecture and AL methods compared to the AL pool. **(A-D)** Comparison of the TFBS content as measured with the Area Under the Receiver Operating Characteristic (AUROC, *y*-axis) for (A, C) human K562 and (B, D) yeast selected sequences across all model architecture/AL method pairs in the (A, B) 3x20k and in the (C, D) 1x60k configurations (*x*-axis). Average AUROC for 15 randomly selected sequence sets compared to the AL pool is represented as a dotted line. **E-H)** Comparison of the di-nucleotide content (*x*-axis) of (E, G) human K562 and (F, H) yeast selected sequences across all NN architecture/AL method pairs in the (E, F) 3x20k and (G, H) 1x60k configurations (y-axis) against the AL pool. Biases for a given di-nucleotide are represented as AUROC heatmaps, red tones indicates an enrichment in the selected sequences compared to the AL pool and blue tones indicates a depletion in the selected sequences compared to the AL pool.

**Supplementary Figure 13:**
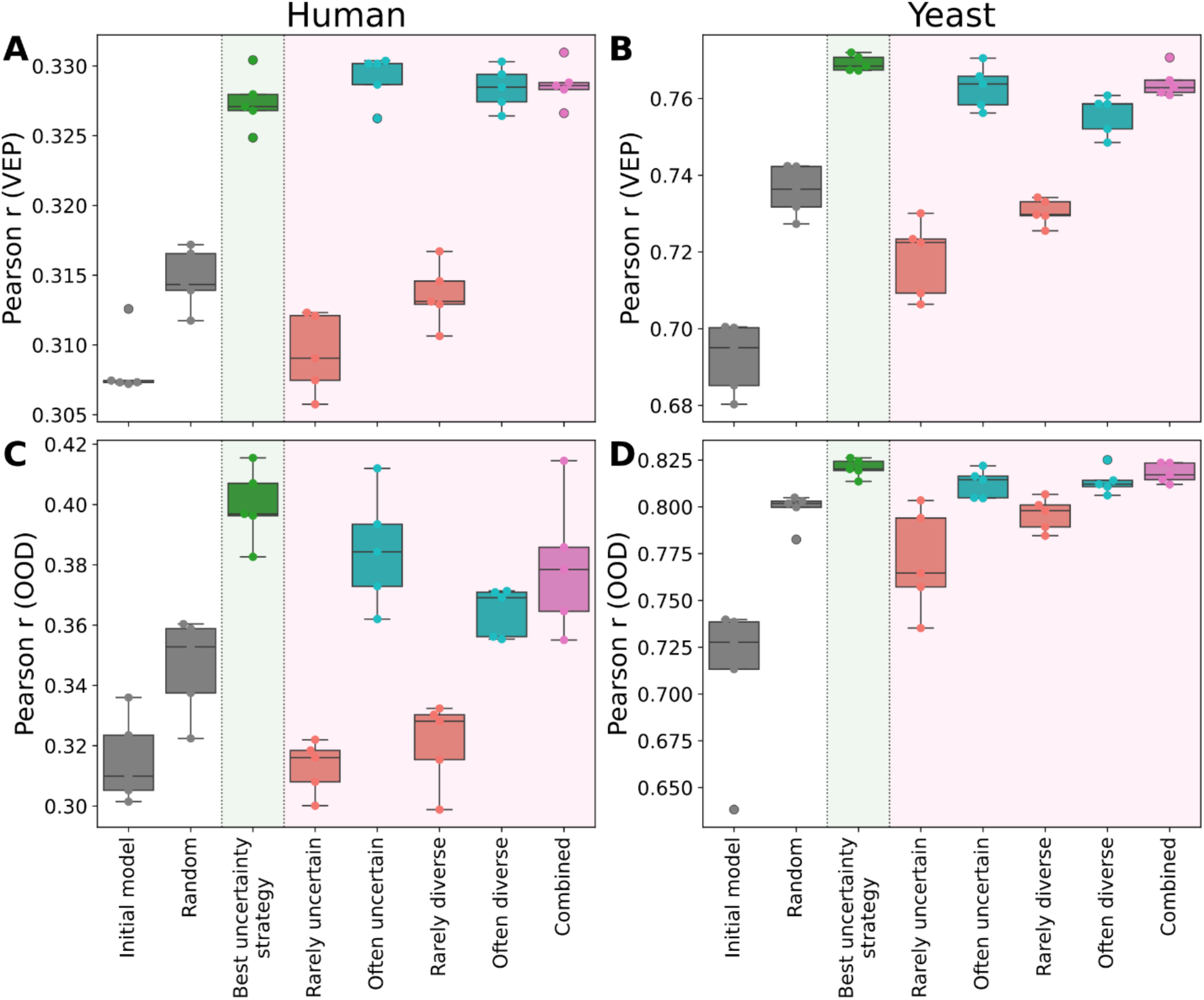
Performance of rarely and often-selected sequences on VEP and OOD test sets, compared to the best individual AL strategy. Performance (Pearson *r*, *y*-axis), of RNN models trained on the rarely- and often-selected sequences by AL (*x*-axis) alongside the base training set in the human (**A, C**) and yeast (**B,D**) datasets, for the VEP (**A-B**) and OOD (**C-D**) test sets. Sequence subsets are broken down for uncertainty- and diversity-based strategies, as well as a “combined” strategy of half most uncertain and half most diverse sequences (pink background). Results are compared to the initial model and random selection (blank background) and to the strongest individual uncertainty-based strategy (green background).

**Supplementary Figure 14:**
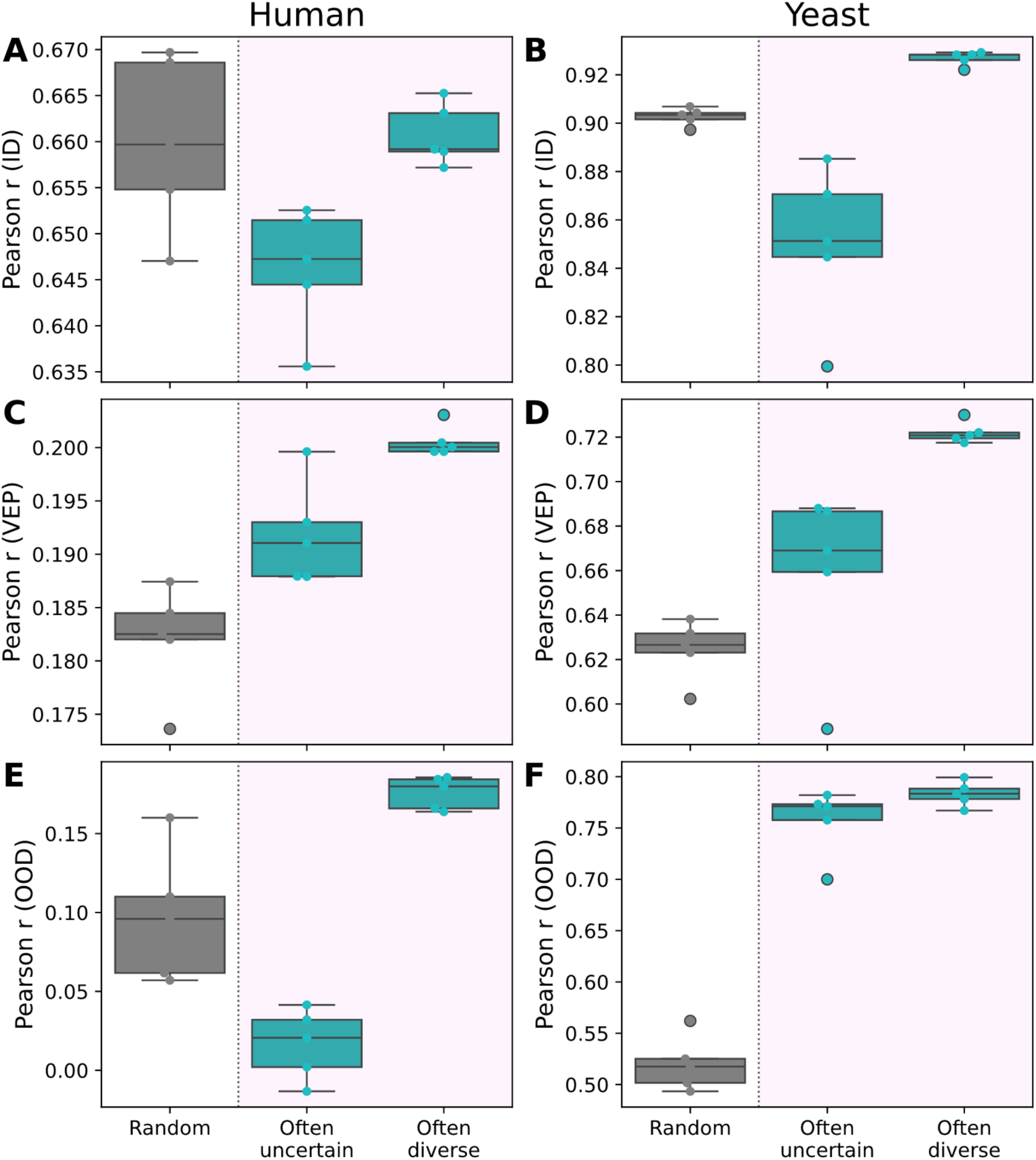
Performance of often-selected sequences as standalone training sets. Performance of RNN models (Pearson *r*, *y*-axes) trained on only the often-selected sequences by uncertainty- and diversity-based AL (i.e. base training set not included; pink background; *x*-axes) in the human (**A, C, E**) and yeast (**B, D, F**) datasets, for the ID (**A-B**), VEP (**C-D**), and OOD (**E-F**) test sets. Results are compared to random selection (blank background).

**Supplementary Figure 15:**
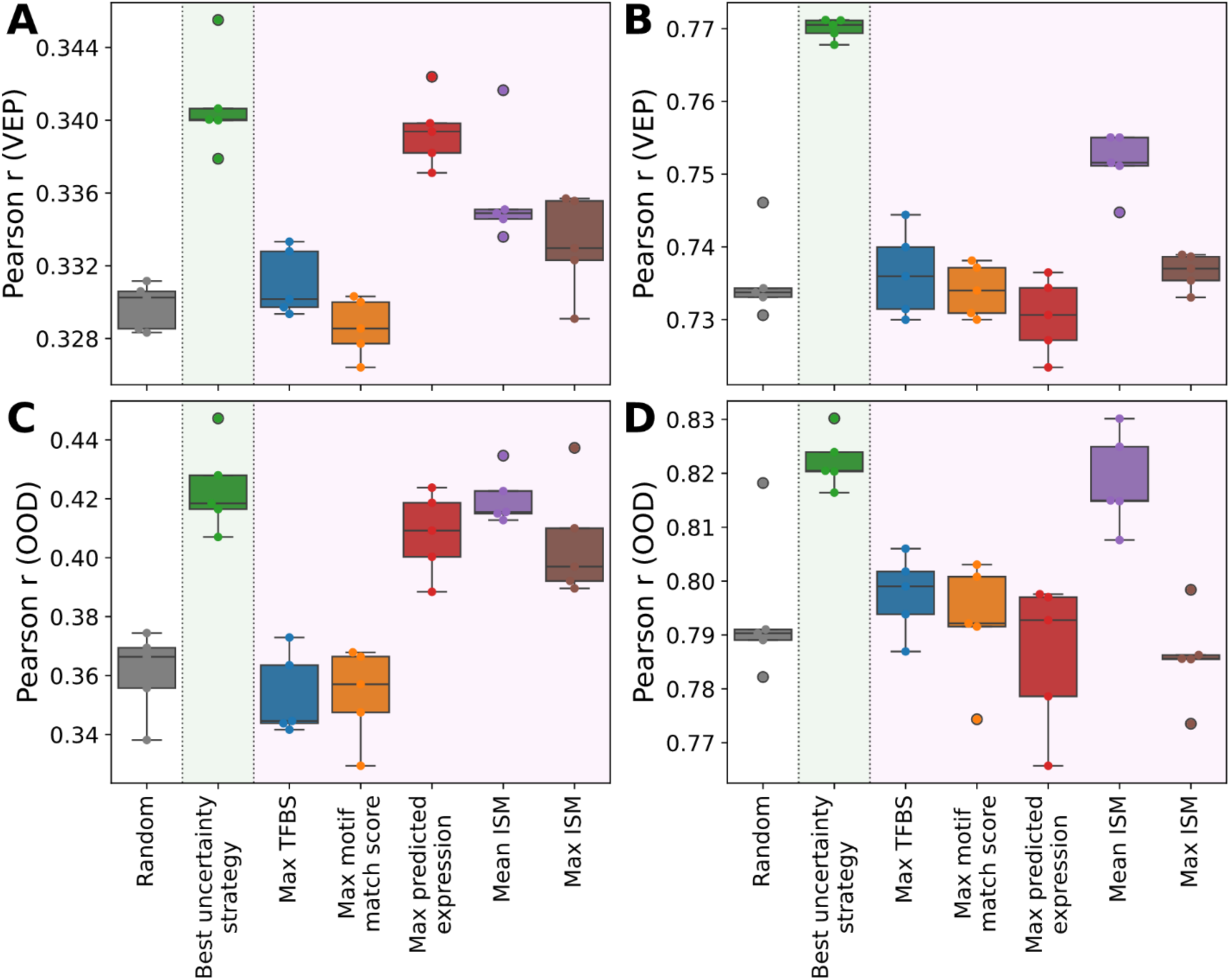
AL produces the most reliable performance gains compared to biologically driven selection strategies. Performance of RNN models (Pearson *r*, *y*-axes) for different sequence selection strategies (*x*-axes), including the best uncertainty-based strategy (green background), random sampling (blank background), and biologically driven sequence acquisition strategies (purple background), on the VEP (**A-B**) and OOD (**C-D**) test sets for human (**A, C**) and yeast (**B, D**) datasets.

## Supplementary Notes

### AL preferentially selects sequences displaying sequence features of high transcriptional activity

We performed a detailed TFBS content analysis for each model architecture/AL method pairs. In human, all fourteen pairs showed AUROC values ranging from 0.54 to 0.64, while random selection showed an AUROC value of 0.50 (**Supplementary Fig. 12A, C**). In yeast, model architecture/AL method pairs-level analysis showed TFBS enrichment for most combinations, and the ensemble-based ones consistently strongly enriched for sequences with high TFBS content compared to the AL pool (AUROC=0.76; **Supplementary Fig. 12B, D**). Hence, in both the human and yeast datasets model architecture and AL methods pairs are preferentially selecting for TFBS rich sequences but to different magnitudes.

We also performed a detailed analysis of the dinucleotide content of selected sequences by all model Architecture and AL method pairs. In human, all pairs, except the model architectures paired with LCMD, tend to modestly enrich for GC dinucleotides and modestly deplete for AT dinucleotides (**Supplementary Fig. 12E, G**). In yeast, the model architectures paired with uncertainty methods showed enrichment for some AT dinucleotides and depletion for some GC dinucleotides (**Supplementary Fig. 12F, H**). The model architectures paired with K-means showed clear enrichment for some GC dinucleotides and depletion for some AT dinucleotides while LCMD containing pairs showed a subtle but similar pattern (**Supplementary Fig. 12F, H**).

Our in-depth analysis indicates that while the often-selected sequences exhibit the most striking enrichment for sequence features of high transcriptional activity like dinucleotide bias and TFBS content, those patterns are also found across most model architecture/AL methods pairs. This indicates that, collectively, AL will preferentially select for sequences containing the DNA features associated with high transcriptional activity. Finally, the above-described patterns are mostly consistent between the 3x20k and the 1x60k configuration, indicating that the different number and size of AL rounds will converge to the selection of sequences with similar properties.

